# Gender imbalances of retraction prevalence among highly cited authors and among all authors

**DOI:** 10.1101/2025.07.08.662297

**Authors:** Stefania Boccia, Antonio Cristiano, Angelo Maria Pezzullo, Jeroen Baas, Guillaume Roberge, John P. A. Ioannidis

## Abstract

**Background:** Scientific retractions remain rare but have become increasingly common. We have previously incorporated retraction data into Scopus-based databases of top-cited (top 2%) scientists to facilitate linkage of retractions with impact metrics at the individual scientist level. Here, we set out to explore whether gender disparities in the likelihood of having retractions exist, both among highly-cited authors and among all authors with ≥5 publications.

**Methods:** We conducted a descriptive cross-sectional bibliometric analysis of a Scopus-based authors database. We used NamSor to assign gender, retaining only results with a confidence >85%. We examined the demographics of scientists with and without retractions among highly cited authors (career-long impact: n = 217,097) and among all other authors (n = 10,361,367). We stratified by publication age, field, country income level, and publication volume, and calculated gender-specific retraction rates and the prevalence ratio (R) of women versus men to have at least one retraction. Multivariable logistic regression were fitted to estimate adjusted associations, reported as odds ratios (OR) and 95% confidence intervals (CI).

**Results:** Gender could be classified for 8,267,888 scientists. Among highly cited authors, 3.3% of men and 2.9% of women had at least one retraction; among all authors, the rate was 0.7% for both genders. Differences varied by field: women’s rates were at least one-third lower than men’s (R < 0.7) in Biology, Biomedical Research, and Psychology (RD<D0.7), but higher (RD>D1.3) in Economics, Engineering, and Information and Communication Technologies. Among highly cited authors, younger cohorts showed increasingly higher rates among men (4.2% men vs. 3.0% women in those starting to publish in 2002–2011; 8.7% men vs. 4.9% women in those starting post-2011). Country-level differences among highly cited authors were pronounced in some countries, as in Pakistan (28.7% men vs. 14.3% women). In multivariable analyses, differences were minimal, if any, for women versus men in having ≥1 retraction (among all authors OR:D0.96, 95% CI: 0.94–0.97) among highly cited authors OR:D0.96, 95% CI: 0.89–1.03), while there were very strong associations of retractions with career age, country income, publication volume, and specific disciplines.

**Conclusion:** Our analysis shows no major gender differences in retraction rates overall, but field-specific differences may exist. Field, career age, country, and publication volume are stronger correlates of retraction. Structural and contextual factors likely drive retraction patterns and warrant further investigation.

## BACKGROUND

Gender disparities in science have been noted in various areas, including recruitment, tenure, funding, authorship, and citation impact. While some of these differences may be narrowing over time, the patterns and changes over time differ among scientific disciplines, environments, and countries(1). Citations play a crucial role as academic influence indicators and contributors to inequalities, particularly among the most-cited scientists, impacting academic career trajectories. A previous study reported that among the 2% top-cited authors for each of 174 science subfields (Science-Metrix classification) of a science-wide author database of standardized citation indicators, men outnumbered women by 1.9-fold (2). Considering 4 publication age cohorts (first publication pre-1992, 1992–2001, 2002–2011, and post-2011), this value decreased from 3.9-fold to 1.4-fold over time.(2)

Recently, the 2% top-cited authors database has been expanded to incorporate retraction data (3), which is often considered a proxy for problematic science. Results show that among 217,097 top-cited scientists in career-long impact and 223,152 in single-year (2023) impact, 7,083 (3.3%) and 8,747 (4.0%), respectively, had at least one retraction. Scientists with retractions had younger publication age, higher self-citation rates, and larger publication volume than those without. No information, however, was available on gender. Notably, in a study examining gender imbalance among retracted biomedical science papers, women comprised 27% of first authors and 24% of last authors, slightly underrepresented compared to estimated general authorship rates of 30–40% for first authors and 25–30% for last authors (4). However, that study did not stratify scientists by publication age cohort or country income level. Importantly, retractions remain an imperfect indicator of problematic science: only a fraction of papers that may warrant retraction for scientific misconduct are ultimately retracted and observed retraction prevalence may partly reflect differences in post-publication scrutiny (5). In this context, accounting for contextual factors may help clarify the role of gender in retraction prevalence and scientific misconduct. In this study, we evaluated gender distribution in retractions among highly cited and all authors worldwide with at least 5 publications, using comprehensive publication and citation data from Scopus and Retraction Watch databases, and tested differences across countries and scientific subfields.

## METHODS

We have generated a comprehensive database of the top 2% most-cited scientists in each of the 174 scientific subfields defined by the Science-Metrix classification (RRID:SCR_024471) (6). This selection was based on a composite citation index, following a methodology similar to our previous studies (7,8). The database also includes scientists who are among the top 100,000 in the composite indicator regardless of their ranking in their primary subfield. The subfields encompass all branches of science, technology, and (bio)medicine, as well as disciplines within the humanities and social sciences. Following a previously established approach (3), we linked Scopus author entries to the Retraction Watch database (RWDB, RRID:SCR_000654), which is the most reliable database of retractions available to date. This linkage allowed us to track the number of retractions associated with each author ID in the Scopus database. Expressions of concern and corrections without retraction, retraction with republication, and retractions where it is explicit that they are due to publisher/journal error rather than author error were excluded.

As in a previous study (2), we employed NamSor (RRID:SCR_023935) (9), a gender-assignment software, to infer the gender of authors in the Scopus database (RRID:SCR_022559). The NamSor algorithm assigns gender based on an author’s first and last name, as well as their country of origin, with a specified confidence level. To determine an author’s country, we used the location of their earliest published paper. We retained only gender assignments with a confidence score above 85%.

We focused on highly cited scientists in the career-long citation impact ranking, including self-citations, and compared them with all Scopus authors with at least five publications. Authors were categorized into four cohorts based on the year of their first publication (i.e., before 1992, 1992-2001, 2002-2011, and after 2011). Within each cohort, authors were classified based on their scientific field, using the 20 major fields defined by the Science-Metrix classification (6). Authors were also categorized by country of the earliest publication and grouped into high-income and other income groups according to the World Bank classification (10). We then calculated the absolute number and proportion of scientists with at least one retracted paper by gender. Specifically, we distinguished men with retractions (M+), men without retractions (M-), women with retractions (F+), and women without retractions (F-). These descriptive analyses were conducted across the four career-age cohorts, countries, and scientific fields.

We examined whether the proportion of retracted authors differed by gender within each subgroup and across different scientific domains. We also calculated the prevalence ratio (R) of women versus men to have a retraction among the top 2% most-cited scientists and among all authors in each subfield. If F+ + F-denotes the total number of women and M+ + M- the total number of men in a given subfield, and F+ and M+ represent the number of women and men with at least one retraction, respectively, then R was calculated as:

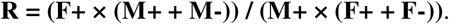

To explore potential drivers of observed differences in retraction rates, we considered overall publication volume. Authors were stratified into tertiles based on their total number of publications, comparing patterns across low-, middle-, and high-publishing groups. We examined the mean and median number of publications in each tertile and across all tertiles and assessed whether differences in retraction rates varied by publication volume.

To estimate the association between gender and the probability of having at least one retracted paper, we fitted multivariable logistic regression models. Corresponding univariable and multivariable models were fitted separately for all authors with at least five publications and for top-cited authors. The models included gender as the main exposure and adjusted for career-age cohort, country income group, citation group (highly cited vs. non-highly cited), scientific field and publication volume expressed as log10(number of papers); the model for all authors also included top-cited status (yes/no). Odds ratios (ORs) and 95% confidence intervals (CIs) were obtained by exponentiating the regression coefficients. In supplementary analyses, we fitted parallel multivariable models that additionally included gender interaction terms with the career-age cohort, income group, publication volume, top-cited status (in the all-authors analysis), and scientific field, to assess whether gender associations varied across strata.

Data was generated and analyzed centrally at Elsevier Research Intelligence. Since Scopus is a subscription database, the full raw data cannot be shared. Accuracy and precision for Scopus have been presented previously (7,8,11). August 2024 dataset version was used for the analyses. The study is reported according to the Strengthening the Reporting of Observational Studies in Epidemiology (STROBE) guidelines (12). Statistical analyses were performed using Python (versio 3.12.3) with numpy (version 2.4.3), pandas (version 3.0.1), statsmodels (version 0.14.6), scipy (version 1.17.1), patsy (version 1.0.2), pyarrow (version 23.0.1) libraries.

## RESULTS

### Overall data on authors and retraction rates in men and women

Out of 10,361,367 authors with at least five full publications, 8,267,888 could be classified for gender (5,295,929 men and 2,971,959 women), while gender was uncertain for 2,093,479 (20.2%). Among the top 2% most-cited authors, gender was confidently assigned for 186,466 individuals: 155,321 men and 31,145 women, while 61,672 (24.9%) had uncertain gender. These entries were excluded from subsequent analyses. Most authors in the overall database came from high-income countries, 6,454,557 of the 10,361,367 authors (62.3%), while 3,482,436 (33.6%) were from middle-or low-income countries. Another 424,374 authors (4.1%) were affiliated with countries that do not have a World Bank income classification, such as small territories, and therefore remained unclassified. Uncertain gender was less frequent in high-income countries where 819,726 of 6,454,557 authors (12.7%) had uncertain gender, whereas in middleD or lowDincome countries 1,003,501 of 3,482,436 authors (28.8%) had uncertain gender.

Retraction patterns by citation level and gender are summarized in **Table 1**. The proportion of authors with at least one retraction was higher among the highly cited scientists, with 7,083 of 217,097 authors (3.3%), than among non-highly cited authors with 72,887 of 10,144,270 (0.7%); this pattern was consistent across genders. Among highly cited authors, 896 of 31,145 women (2.9%) and 4,752 of 155,321 men (3.1%) had at least one retraction. Among non-highly cited authors, 19,693 of 2,940,814 women (0.7%) and 33,816 of 5,140,608 men (0.7%) had at least one retraction.. In nonDhighDincome countries, 1,628 of 22,181 highly cited authors (7.3%) and 49,645 of 3,460,255 nonDhighly cited authors (1.4%) had at least one retraction, compared with 5,439 of 193,694 highly cited authors (2.8%) and 23,030 of 6,260,863 nonDhighly cited authors (0.4%) in highDincome countries. After stratifying by income, men had slightly higher retraction rates than women in both country groups.

**Table 1.** Number and percentage of authors with at least one retraction by gender and income level, stratified by highly cited and non-highly cited authors.

|  | All citation group |  | Highly cited authors |  | Non-highly cited authors |  |
| --- | --- | --- | --- | --- | --- | --- |
|  | All authors, n | With retraction, n (%) | All authors, n | With retraction, n (%) | All authors, n | With retraction, n (%) |
| **All income levels** |  |  |  |  |  |  |
| Overall | 10,361,367 | 79,970 (0.8) | 217,097 | 7083 (3.3) | 10,144,270 | 72,887 (0.7) |
| Women | 2,971,959 | 20,589 (0.7) | 31,145 | 896 (2.9) | 2,940,814 | 19,693 (0.7) |
| Men | 5,295,929 | 38,568 (0.7) | 155,321 | 4752 (3.1) | 5,140,608 | 33,816 (0.7) |
| **High income levels** |  |  |  |  |  |  |
| Overall | 6,454,557 | 28,469 (0.4) | 193,694 | 5439 (2.8) | 6,260,863 | 23,030 (0.4) |
| Women | 1,931,003 | 72,55 (0.4) | 28,009 | 696 (2.5) | 1,902,994 | 6559 (0.3) |
| Men | 3,700,367 | 17,203 (0.5) | 141,680 | 3865 (2.7) | 3,558,687 | 13,338 (0.4) |
| **Other income levels** |  |  |  |  |  |  |
| Overall | 3,482,436 | 51,273 (1.5) | 22,181 | 1628 (7.3) | 3,460,255 | 49,645 (1.4) |
| Women | 1,002,636 | 13,277 (1.3) | 2981 | 198 (6.6) | 999,655 | 13,079 (1.3) |
| Men | 1,475,514 | 21,237 (1.4) | 12,724 | 876 (6.9) | 1,462,790 | 20,361 (1.4) |

Highly cited authors had far more publications than non-highly cited ones, with a median of 163 vs. 11 for all authors regardless of gender (data not shown), and the same pattern across income strata (**Supplementary Table 1, Additional File 1**). Men had modestly more publications than women (median 14 vs. 11), a pattern also observed across strata. Authors from non-high-income countries who were highly cited had slightly more publications than their high-income counterparts (median 190 vs. 160, data not shown). In non-high-income countries, median publication count was slightly higher in women than men (189 versus 176), whereas the opposite was true in high-income countries (162 vs. 137; Supplementary Table 1, Additional File 1). The majority of authors with at least one retraction, 52,967 of 79,970 (66.2%), were in the first tertile of publication counts (Supplementary Table 2, Additional File 1).

### Retraction rates in men and women across scientific fields

**Table 2** presents retraction rates across scientific fields by gender for all authors, and separately for highly cited and not highly cited authors. While gender differences were generally small, some patterns emerged. Excluding fields with fewer than 50 authors with retractions and thus high uncertainty, women had retraction rates at least one-third lower than men (R<0.7) in Biology, Biomedical Research, and Psychology & Cognitive Sciences. Conversely, in Economics & Business, Engineering, and Information & Communication Technologies, retractions appeared more frequently among women (R>1.3). R values for non-highly cited authors closely matched those for all authors. Highly cited authors, however, showed different patterns, with highest R values in Mathematics & Statistics (R=3.1) and Engineering (R=1.8), indicating higher retraction likelihood among women. In contrast, lower R values among highly cited authors were in Biomedical Research (R=0.6), Built Environment & Design (R=0.6), and Economics & Business (R=0.7), suggesting lower retraction rates among women.

**Table 2.** Overall number and percentage of authors by gender with at least one retraction, according to subfield, along with the ratio of retractions among women over men (R), stratified by highly cited and non-highly cited authors.

| Field | All authors |  |  | Highly cited authors |  |  | Non-highly cited authors |  |  |
| --- | --- | --- | --- | --- | --- | --- | --- | --- | --- |
|  | Women, n (%) | Men, n (%) | R | Women, n (%) | Men, n (%) | R | Women, n (%) | Men, n (%) | R |
| Agriculture, Fisheries & Forestry | 397 (0.3) | 887 (0.5) | 0.68 | 15 (1.4) | 60 (1.1) | 1.19 | 382 (0.3) | 827 (0.5) | 0.69 |
| Biology | 600 (0.5) | 1390 (0.7) | 0.65 | 33 (2.7) | 151 (2.3) | 1.17 | 567 (0.4) | 1239 (0.7) | 0.67 |
| Biomedical Research | 2215 (0.7) | 3,696 (1.0) | 0.65 | 92 (3.2) | 609 (5.1) | 0.64 | 2123 (0.6) | 3087 (0.9) | 0.72 |
| Built Environment & Design | 43 (0.3) | 132 (0.4) | 0.79 | 3 (1.6) | 21 (2.5) | 0.65 | 40 (0.3) | 111 (0.3) | 0.87 |
| Chemistry | 902 (0.4) | 1842 (0.5) | 0.87 | 52 (3.1) | 284 (2.7) | 1.16 | 850 (0.4) | 1558 (0.4) | 0.95 |
| Clinical Medicine | 10,879 (1.0) | 18,647 (1.2) | 0.85 | 441 (4.0) | 2289 (4.7) | 0.85 | 10,438 (1.0) | 16,358 (1.1) | 0.91 |
| Communication & Textual Studies | 13 (0.1) | 24 (0.1) | 0.62 | 1 (0.3) | 1 (0.2) | 1.63 | 12 (0.1) | 23 (0.1) | 0.59 |
| Earth & Environmental Sciences | 480 (0.5) | 913 (0.5) | 1.05 | 16 (2.0) | 91 (1.6) | 1.21 | 464 (0.5) | 822 (0.5) | 1.11 |
| Economics & Business | 343 (0.6) | 509 (0.4) | 1.41 | 6 (1.0) | 44 (1.4) | 0.68 | 337 (0.6) | 465 (0.4) | 1.49 |
| Enabling & Strategic Technologies | 1100 (0.6) | 2624 (0.5) | 1.14 | 71 (3.4) | 384 (3.1) | 1.10 | 1029 (0.6) | 2240 (0.5) | 1.23 |
| Engineering | 899 (0.7) | 1967 (0.4) | 1.58 | 52 (3.6) | 247 (2.0) | 1.78 | 847 (0.6) | 1720 (0.4) | 1.68 |
| Historical Studies | 7 (0.0) | 28 (0.1) | 0.45 | 0 (0.0) | 0 (0.0) | - | 7 (0.0) | 28 (0.1) | 0.45 |
| Information & Communication Technologies | 1580 (1.1) | 3345 (0.7) | 1.43 | 22 (1.4) | 186 (1.7) | 0.81 | 1558 (1.1) | 3159 (0.7) | 1.48 |
| Mathematics & Statistics | 100 (0.4) | 336 (0.4) | 0.99 | 6 (4.0) | 28 (1.3) | 3.06 | 94 (0.4) | 308 (0.4) | 0.99 |
| Philosophy & Theology | 8 (0.1) | 13 (0.1) | 1.80 | 0 (0.0) | 0 (0.0) | - | 8 (0.1) | 13 (0.1) | 1.79 |
| Physics & Astronomy | 461 (0.3) | 1578 (0.3) | 1.05 | 30 (2.3) | 252 (1.6) | 1.43 | 431 (0.3) | 1326 (0.3) | 1.14 |
| Psychology & Cognitive Sciences | 190 (0.3) | 282 (0.5) | 0.62 | 24 (2.3) | 65 (2.6) | 0.89 | 166 (0.3) | 217 (0.4) | 0.68 |
| Public Health & Health Services | 205 (0.2) | 151 (0.3) | 0.71 | 28 (1.6) | 27 (1.6) | 0.96 | 177 (0.2) | 124 (0.2) | 0.73 |
| Social Sciences | 166 (0.2) | 204 (0.2) | 0.97 | 4 (0.3) | 13 (0.4) | 0.66 | 162 (0.2) | 191 (0.2) | 1.00 |
| Visual & Performing Arts | 1 (0.0) | 0 (0.0) | N/A | 0 (0.0) | 0 (0.0) | - | 1 (0.0) | 0 (0.0) | N/A |
| All fields | 20,589 (0.7) | 38,568 (0.7) | 0.95 | 896 (2.9) | 4752 (3.1) | 0.94 | 19,693 (0.7) | 33,816 (0.7) | 1.02 |

N/A, not applicable.

### Retraction rates in men and women across publication age cohorts

Overall, men consistently had slightly higher retraction rates than women across all different publication age cohorts (**Figure 1a**), although the difference was small in percentage terms. Among highly cited authors (**Figure 1b**), men and women had similar retraction rates in the two older cohorts, while men had higher rates than women in the 2002–2011 cohort (470 of 11,184 men, 4.2% *vs*. 46 of 1,534 women, 3.0%) and ≥2012 cohorts (2,604 of 29,928 men, 8.7% *vs*. 129 of 2,641 women, 4.9%). Supplementary Tables 3–S break down retraction rates by cohort for highly cited, non-highly cited, and all authors. Among highly cited scientists (**Supplementary Table 3, Additional File 1**), retraction rates for men and women were close in earlier cohorts but diverged across specific fields. In Clinical Medicine, the gender difference widened: women had retraction rates of 3.9% (n=255) pre-1992 and 4.6% (n=156) in 1992–2001, dropping to 3.0% (n=30) in 2002–2011 and 0% (n=0) post-2011. In contrast, men’s rates rose over time from 4.5% (n=1,637) to 5.4% (n=503), 5.5% (n=138) and 6.5% (n=11). In Enabling & Strategic Technologies, both genders showed rising retraction rates. Among women, rates increased from 2.6% (n=16) to 3.8% (n=28), 3.6% (n=24), and 6.3% (n=3). In Information & Communication Technologies, women had higher rates in the first cohort (2.1%; n=10), followed by a decline to 0.8% (n=36), 1.6% (n=7) and 0%. Men’s rates, by contrast, rose steadily from 0.8% (n=36) to 9.1% (n=20). In Engineering, rates among women increased from 2.2% (n=11) to 4.6% (n=17) in the first two cohorts, while among men rose from 1.2% (n=85) to 3.5% (n=66). Most other fields had too small numbers of highly cited authors in the youngest (post-2011) cohorts to make any meaningful inferences.

**Figure 1.**
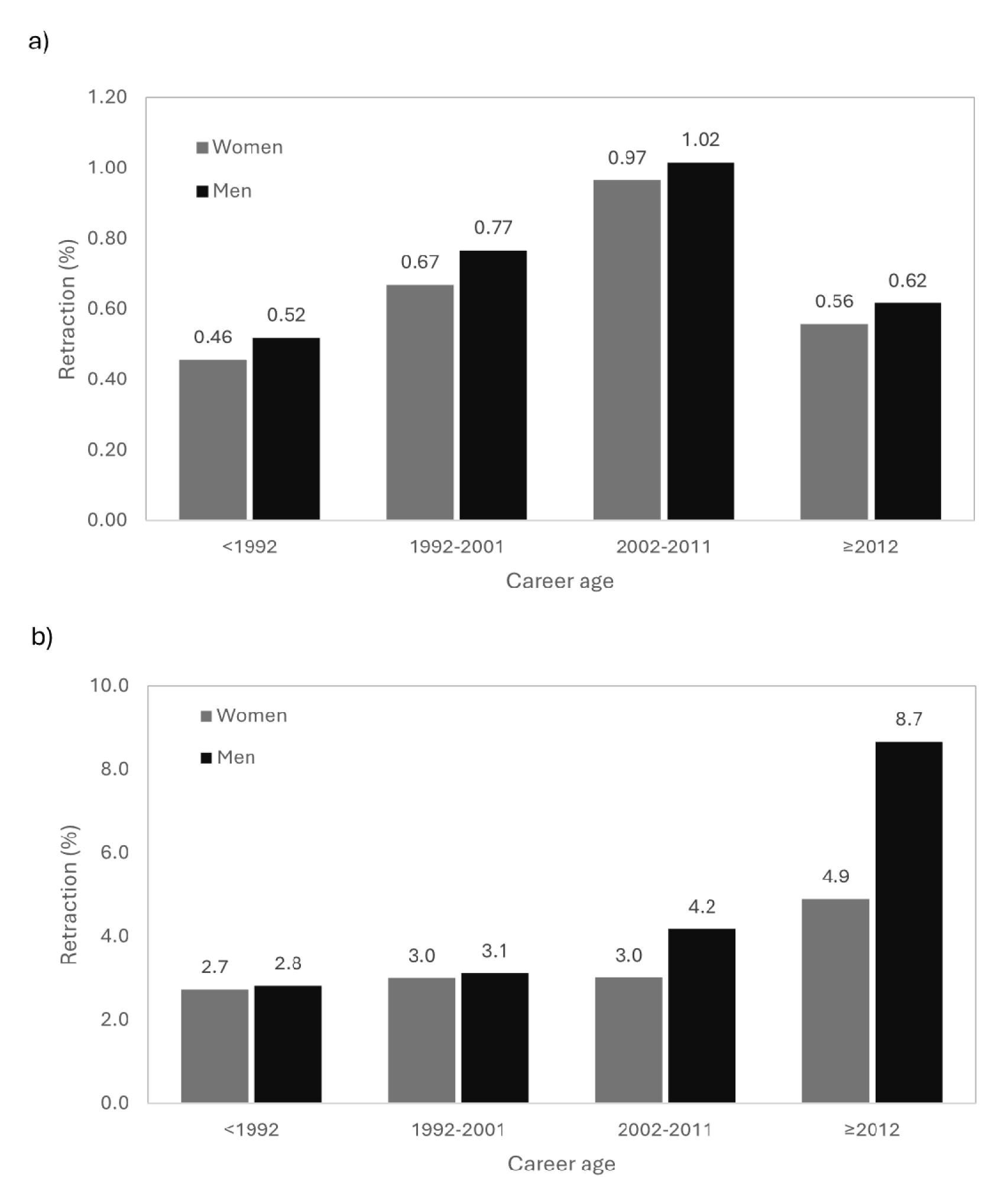
Proportion of authors with at least one retraction by gender and career age (year of first publication) across all citation groups (a) and among highly cited scientists (b).

Among non-highly cited authors (**Supplementary Table 4, Additional File 1**), retraction rates increased over time for both genders. In Clinical Medicine, where absolute numbers were highest, the gender difference remained roughly constant: men’s rates rose from 0.5% (n=2,111) to 1.2% (n=5,081), women’s from 0.4% (n=623) to 1.0% (n=4,113). In Engineering, the gender difference narrowed: in 1992–2001, rates were 0.7% (n=102) for women and 0.4% (n=271) for men; in 2002–2011, 1.1% (n=539) vs. 0.7% (n=959); and in postD2011, 0.3% (n=190) vs. 0.2% (n=388). In contrast, Information & Communication Technologies showed a persistent difference with higher rates in women, especially in 2002–2011 (1.8% [n=970] vs. 1.1% [n=1,721]), dissimilar to patterns in the highly cited group. When all fields were combined, there was no gender difference across cohorts among non-highly cited authors. When all authors were combined (**Supplementary Table 5, Additional File 1**), trends closely mirrored those in the non-highly cited group, with no substantial differences in rates or gender patterns by cohort.

### Retraction rates in men and women across different countries

Among the 40 countries with the highest number of authors (**Table 3**), some countries showed a comparatively high proportion of highly cited women with retractions, even though the denominators were relatively small. In Pakistan, 2 of 14 women (14.3%) and 52 of 181 men (28.7%) had at least one retraction. In Egypt, 4 of 29 women (13.8%) and 37 of 411 men (9.0%); in Iran, 5 of 54 women (9.3%) and 103 of 830 men (12.4%); in China, 148 of 1,821 women (8.1%) and 325 of 4,204 men (7.7%); in India, 16 of 242 women (6.6%) and 195 of 2,116 men (9.2%); in Taiwan, 14 of 217 women (6.5%) and 27 of 643 men (4.2%); in Italy, 62 of 1,082 women (5.7%) and 172 of 4,687 men (3.7%); and in the Czech Republic, 3 of 55 women (5.5%) and 14 of 428 men (3.3%). The relative gender difference was particularly notable in Pakistan, and similar albeit attenuated patterns were observed in Iran and India. Conversely, women had higher retraction rates than men in Italy, Taiwan, the Czech Republic, and Egypt, but the absolute numbers are low and the differences modest.

**Table 3.** Number of authors with at least one retraction by gender in the 40 countries with the highest number of authors with ≥5 publications.

| Country | All authors |  | Highly cited authors |  | Non-highly cited authors |  |
| --- | --- | --- | --- | --- | --- | --- |
|  | Women, n (%) | Men, n (%) | Women, n (%) | Men, n (%) | Women, n (%) | Men, n (%) |
| United States | 2520 (0.4) | 5634 (0.5) | 340 (2.5) | 1688 (2.7) | 2180 (0.4) | 3946 (0.3) |
| China | 10,296 (2.1) | 12,777 (2.2) | 148 (8.1) | 325 (7.7) | 10,148 (2.1) | 12,452 (2.1) |
| Japan | 296 (0.4) | 1987 (0.5) | 8 (2.3) | 238 (4.1) | 288 (0.3) | 1749 (0.4) |
| Germany | 340 (0.3) | 1055 (0.3) | 33 (3.2) | 259 (2.7) | 307 (0.3) | 796 (0.3) |
| United Kingdom | 360 (0.3) | 928 (0.4) | 60 (1.9) | 284 (1.9) | 300 (0.2) | 644 (0.3) |
| India | 909 (1.1) | 2978 (1.6) | 16 (6.6) | 195 (9.2) | 893 (1.1) | 2783 (1.5) |
| France | 348 (0.3) | 628 (0.4) | 12 (1.3) | 111 (2.2) | 336 (0.3) | 517 (0.3) |
| Italy | 847 (0.7) | 984 (0.6) | 62 (5.7) | 172 (3.7) | 785 (0.6) | 812 (0.5) |
| Russia | 181 (0.3) | 219 (0.2) | 1 (1.4) | 11 (1.3) | 180 (0.3) | 208 (0.2) |
| Canada | 267 (0.3) | 557 (0.4) | 34 (2.2) | 167 (2.6) | 233 (0.3) | 390 (0.3) |
| Spain | 278 (0.3) | 481 (0.4) | 4 (0.7) | 71 (2.8) | 274 (0.3) | 410 (0.4) |
| South Korea | 188 (0.6) | 844 (0.8) | 6 (3.6) | 56 (4.6) | 182 (0.6) | 788 (0.8) |
| Brazil | 209 (0.3) | 294 (0.3) | 4 (2.5) | 18 (2.4) | 205 (0.2) | 276 (0.3) |
| Australia | 216 (0.3) | 439 (0.5) | 27 (2.1) | 124 (2.4) | 189 (0.3) | 315 (0.4) |
| Netherlands | 124 (0.3) | 271 (0.4) | 17 (3.0) | 84 (2.4) | 107 (0.2) | 187 (0.3) |
| Poland | 98 (0.2) | 108 (0.2) | 6 (2.7) | 13 (1.6) | 92 (0.2) | 95 (0.2) |
| Turkey | 169 (0.5) | 440 (0.7) | 2 (1.4) | 22 (2.6) | 167 (0.5) | 418 (0.7) |
| Switzerland | 85 (0.3) | 253 (0.4) | 2 (0.5) | 72 (2.7) | 83 (0.3) | 181 (0.3) |
| Taiwan | 201 (0.8) | 285 (0.7) | 14 (6.5) | 27 (4.2) | 187 (0.8) | 258 (0.7) |
| Sweden | 102 (0.3) | 238 (0.5) | 8 (1.7) | 65 (2.6) | 94 (0.3) | 173 (0.3) |
| Iran | 360 (1.3) | 1184 (2.1) | 5 (9.3) | 103 (12.4) | 355 (1.2) | 1081 (2.0) |
| Belgium | 49 (0.2) | 125 (0.3) | 7 (3.0) | 32 (2.1) | 42 (0.2) | 93 (0.2) |
| Mexico | 44 (0.2) | 95 (0.3) | 1 (2.2) | 5 (1.7) | 43 (0.2) | 90 (0.3) |
| Austria | 34 (0.2) | 99 (0.3) | 1 (0.7) | 19 (1.7) | 33 (0.2) | 80 (0.2) |
| Denmark | 41 (0.2) | 123 (0.4) | 6 (2.1) | 37 (2.3) | 35 (0.2) | 86 (0.3) |
| Indonesia | 31 (0.2) | 82 (0.4) | 0 (0.0) | 2 (8.0) | 31 (0.2) | 80 (0.3) |
| Israel | 74 (0.4) | 136 (0.5) | 1 (0.4) | 27 (1.7) | 73 (0.4) | 109 (0.4) |
| Czech Republic | 67 (0.4) | 138 (0.5) | 3 (5.5) | 14 (3.3) | 64 (0.4) | 124 (0.4) |
| Malaysia | 144 (1.0) | 300 (1.2) | 3 (7.1) | 27 (11.1) | 141 (1.0) | 273 (1.1) |
| Finland | 30 (0.2) | 63 (0.2) | 5 (1.6) | 25 (2.6) | 25 (0.1) | 38 (0.2) |
| Egypt | 178 (1.7) | 551 (1.9) | 4 (13.8) | 37 (9.0) | 174 (1.7) | 514 (1.8) |
| Portugal | 47 (0.2) | 60 (0.3) | 6 (4.6) | 8 (1.8) | 41 (0.2) | 52 (0.2) |
| Ukraine | 18 (0.1) | 29 (0.2) | 0 (0.0) | 1 (1.2) | 18 (0.1) | 28 (0.2) |
| Greece | 57 (0.4) | 144 (0.5) | 3 (2.6) | 23 (2.7) | 54 (0.4) | 121 (0.5) |
| Norway | 45 (0.3) | 76 (0.3) | 0 (0.0) | 21 (2.0) | 45 (0.3) | 55 (0.2) |
| Argentina | 61 (0.3) | 57 (0.3) | 0 (0.0) | 5 (2.8) | 61 (0.3) | 52 (0.3) |
| Thailand | 52 (0.4) | 69 (0.5) | 0 (0.0) | 6 (4.3) | 52 (0.4) | 63 (0.5) |
| Pakistan | 201 (2.0) | 920 (3.8) | 2 (14.3) | 52 (28.7) | 199 (2.0) | 868 (3.6) |
| Romania | 71 (0.4) | 84 (0.5) | 1 (2.6) | 5 (4.1) | 70 (0.4) | 79 (0.5) |
| South Africa | 48 (0.4) | 64 (0.4) | 5 (3.6) | 8 (1.4) | 43 (0.4) | 56 (0.3) |

In most other countries the differences were negligible. For example, in the United States 340 of 13,489 women (2.5%) and 1,688 of 61,731 men (2.7%) had at least one retraction; in the United Kingdom 60 of 3,112 women (1.9%) versus 284 of 14,717 men (1.9%); and in Canada 34 of 1,555 women (2.2%) and 167 of 6,535 men (2.6%).

Among all authors, regardless of citation level, gender differences remained limited. Women had lower retraction rates than men in India, with 909 of 79,242 women (1.1%) versus 2,978 of 186,143 men (1.6%), in Pakistan, with 201 of 10,069 women (2.0%) versus 920 of 24,339 men (3.8%), and in Iran, with 360 of 28,483 women (1.3%) versus 1,184 of 55,983 men (2.1%). By contrast, slightly higher rates in women were observed in Italy, where 847 of 126,198 women (0.7%) had at least one retraction compared with 984 of 159,951 men (0.6%), and in South Africa, where 48 of 11,946 women (0.4%) versus 64 of 17,902 men (0.4%). In most countries, gender differences remained small in this broader analysis. Patterns for nonDhighly cited authors were similar (not shown).

### Logistic regression analyses

We report on univariable and multivariable logistic regression analyses in **Table 4**. In multivariable analyses, women had minimally lower odds of having at least one retraction than men among all authors (OR 0.96, 95% CI 0.94–0.97), while the corresponding estimate among top-cited authors was similarly small and not clearly different from 1 (OR 0.96, 95% CI 0.89–1.03). By contrast, very strong and consistent associations were seen for career age, country income level, and publication volume. Compared with authors whose first publication predated 1992, all three younger cohorts had higher odds of retraction, especially those starting in 2002–2011 and ≥2012 (all authors OR 3.14 95% CI 3.06–3.22 and 2.91 95% CI 2.83–2.99; highly-cited authors OR 1.87 95% CI 1.73–2.03 and 4.67 95% CI 3.89–5.62, respectively). Authors from non-high-income countries also had markedly higher odds than those from high-income countries (all authors OR 4.17, 95% CI 4.10–4.24; highly-cited authors OR 2.56, 95% CI 2.39–2.75), and publication volume was a strong predictor in both analyses (per 1-unit increase in log10 papers: OR 8.69, 95% CI 8.54–8.85, and OR 11.36, 95% CI 10.40–12.40, respectively). Highly-cited status itself was associated with only a modest increase in odds in all authors model (OR 1.40, 95% CI 1.36–1.44). Across scientific fields, adjusted odds were lower than in Clinical Medicine for almost all fields, with Biomedical Research among highly-cited authors the main exception (OR 1.36 95% CI 1.25–1.47). Multivariable models including interaction terms (**Supplementary Table 6, Additional File 1**), suggested lower retraction risk among all authors for women in the youngest cohort, in non-high income countries, in Biology and Biomedical Research and higher retraction risk for women in some other scientific fields (Earth & Environmental Sciences, Economics & Business, Enabling & Strategic Technologies, Engineering, Information & Communication Technologies, Physics & Astronomy).

**Table 4.** Univariable and multivariable odds ratios (OR) and 95% confidence intervals (CI) from logistic regression models estimating the probability of having >=1 retraction, among all authors and top-cited authors. Publication volume is log10-transformed (number of papers). Reference categories: Men; <1992; High income level; No top-cited status; Clinical Medicine.

|  | All authors |  |  |  | Top-cited authors |  |  |  |
| --- | --- | --- | --- | --- | --- | --- | --- | --- |
|  | n | n retracted | Univariable OR (95% CI) | Multivariable OR (95% CI) | n | n retracted | Univariable OR (95% CI) | Multivariable OR (95% CI) |
| *Gender* |  |  |  |  |  |  |  |  |
| Men | 5,295,929 | 38,568 | Ref | Ref | 155,321 | 4,752 | Ref | Ref |
| Women | 2,971,959 | 20,589 | 0.95 (0.93-0.97) | 0.96 (0.94-0.97) | 31,145 | 896 | 0.94 (0.87-1.01) | 0.96 (0.89-1.03) |
| Unknown | 2,093,479 | 20,813 | 1.37 (1.35-1.39) | 1.24 (1.22-1.26) | 30,631 | 1,435 | 1.56 (1.47-1.65) | 1.16 (1.08-1.23) |
| *Age career (year of first publication)* |  |  |  |  |  |  |  |  |
| <1992 | 2,306,544 | 9,911 | Ref | Ref | 137,364 | 4,092 | Ref | Ref |
| 1992-2001 | 1,529,389 | 12,200 | 1.86 (1.81-1.91) | 1.86 (1.81-1.91) | 54,909 | 1,879 | 1.15 (1.09-1.22) | 1.31 (1.23-1.39) |
| 2002-2011 | 2,889,339 | 33,469 | 2.72 (2.66-2.78) | 3.14 (3.06-3.22) | 23,004 | 964 | 1.42 (1.33-1.53) | 1.87 (1.73-2.03) |
| $\geq 2012$ | 3,636,095 | 24,390 | 1.56 (1.53-1.60) | 2.91 (2.83-2.99) | 1,820 | 148 | 2.88 (2.43-3.42) | 4.67 (3.89-5.62) |
| *Income level* |  |  |  |  |  |  |  |  |
| High income level | 6,454,557 | 28,469 | Ref | Ref | 193,694 | 5,439 | Ref | Ref |
| All other income levels | 3,482,436 | 51,273 | 3.37 (3.32-3.42) | 4.17 (4.10-4.24) | 22,181 | 1,628 | 2.74 (2.59-2.90) | 2.56 (2.39-2.75) |
| Unknown | 424,374 | 228 | 0.12 (0.11-0.14) | 0.31 (0.27-0.35) | 1,222 | 16 | 0.46 (0.28-0.75) | 0.59 (0.35-0.97) |
| *Publication volume (log10 papers)* |  |  |  |  |  |  |  |  |
| Per +1 unit | 10,361,367 | 79,970 | 5.65 (5.58-5.72) | 8.69 (8.54-8.85) | 217,097 | 7,083 | 12.00 (11.08-13.00) | 11.36 (10.40-12.40) |
|  | n | n retracted | Univariable OR (95% CI) | Multivariable OR (95% CI) | n | n retracted | Univariable OR (95% CI) | Multivariable OR (95% CI) |
| *Highly-cited status* |  |  |  |  |  |  |  |  |
| No | 10,144,270 | 72,887 | Ref | Ref | 0 | 0 |  |  |
| Yes | 217,097 | 7,083 | 4.66 (4.55-4.78) | 1.40 (1.36-1.44) | 217,097 | 7,083 |  |  |
| *Scientific field* |  |  |  |  |  |  |  |  |
| Clinical Medicine | 3,280,471 | 39,335 | Ref | Ref | 67,839 | 3,249 | Ref | Ref |
| Agriculture, Fisheries & Forestry | 368,413 | 1,663 | 0.37 (0.36-0.39) | 0.28 (0.26-0.29) | 7,265 | 99 | 0.27 (0.22-0.34) | 0.38 (0.31-0.46) |
| Biology | 391,318 | 2,489 | 0.53 (0.51-0.55) | 0.42 (0.40-0.44) | 8,656 | 222 | 0.52 (0.46-0.60) | 0.83 (0.72-0.96) |
| Biomedical Research | 831,462 | 7,298 | 0.73 (0.71-0.75) | 0.82 (0.80-0.84) | 16,898 | 846 | 1.05 (0.97-1.13) | 1.36 (1.25-1.47) |
| Built Environment & Design | 59,638 | 219 | 0.30 (0.27-0.35) | 0.29 (0.26-0.34) | 1,243 | 34 | 0.56 (0.40-0.79) | 0.70 (0.49-0.99) |
| Chemistry | 723,834 | 3,650 | 0.42 (0.40-0.43) | 0.30 (0.29-0.31) | 14,911 | 462 | 0.64 (0.58-0.70) | 0.50 (0.45-0.56) |
| Communication & Textual Studies | 54,770 | 42 | 0.06 (0.05-0.09) | 0.12 (0.09-0.17) | 1,074 | 2 | 0.04 (<0.01-0.15) | 0.14 (0.04-0.57) |
| Earth & Environmental Sciences | 351,435 | 1,943 | 0.46 (0.44-0.48) | 0.34 (0.32-0.35) | 7,388 | 157 | 0.43 (0.37-0.51) | 0.50 (0.43-0.59) |
| Economics & Business | 197,907 | 1,170 | 0.49 (0.46-0.52) | 0.58 (0.55-0.62) | 4,137 | 59 | 0.29 (0.22-0.37) | 0.60 (0.46-0.78) |
| Enabling & Strategic Technologies | 926,899 | 5,402 | 0.48 (0.47-0.50) | 0.31 (0.30-0.32) | 18,317 | 654 | 0.74 (0.68-0.80) | 0.51 (0.46-0.56) |
| Engineering | 800,260 | 4,360 | 0.45 (0.44-0.47) | 0.32 (0.31-0.33) | 17,118 | 432 | 0.51 (0.46-0.57) | 0.45 (0.41-0.50) |
| Historical Studies | 52,202 | 37 | 0.06 (0.04-0.08) | 0.12 (0.08-0.16) | 1,081 | 0 | — | — |
| Information & Communication | 757,752 | 7,671 | 0.84 (0.82-0.86) | 0.58 (0.56-0.59) | 15,087 | 275 | 0.37 (0.33-0.42) | 0.31 (0.28-0.36) |
|  | n | n<br>retracted | Univariable OR<br>(95% CI) | Multivariable<br>OR (95% CI) | n | n<br>retracted | Univariable OR<br>(95% CI) | Multivariable<br>OR (95% CI) |
| Technologies |  |  |  |  |  |  |  |  |
| Mathematics & Statistics | 126,690 | 570 | 0.37 (0.34-0.40) | 0.32 (0.29-0.35) | 2,693 | 48 | 0.36 (0.27-0.48) | 0.47 (0.35-0.63) |
| Philosophy & Theology | 28,024 | 22 | 0.06 (0.04-0.10) | 0.13 (0.09-0.20) | 523 | 0 | — | — |
| Physics & Astronomy | 851,450 | 2,706 | 0.26 (0.25-0.27) | 0.16 (0.15-0.16) | 19,980 | 361 | 0.37 (0.33-0.41) | 0.32 (0.28-0.35) |
| Psychology & Cognitive Sciences | 127,443 | 529 | 0.34 (0.32-0.37) | 0.55 (0.51-0.60) | 3,871 | 98 | 0.52 (0.42-0.63) | 0.90 (0.73-1.10) |
| Public Health & Health Services | 194,989 | 425 | 0.18 (0.16-0.20) | 0.25 (0.23-0.27) | 3,841 | 65 | 0.34 (0.27-0.44) | 0.51 (0.40-0.66) |
| Social Sciences | 228,291 | 438 | 0.16 (0.14-0.17) | 0.26 (0.23-0.28) | 5,063 | 20 | 0.08 (0.05-0.12) | 0.22 (0.14-0.34) |
| Visual & Performing Arts | 7,277 | 1 | 0.01 (<0.01-0.08) | 0.04 (<0.01-0.25) | 112 | 0 | — | — |

## DISCUSSION

Our analyses show that, overall, retraction rates are similar in women and men among all authors and among highly cited authors. A similar picture emerged after adjusting for career age, country income level, publication volume, and scientific field. These variables were far stronger correlates of the risk of retraction than gender. The retraction rates correlated strongly with younger career age, provenance from a country with no high income, a higher publication volume, and certain scientific fields than others. When examining scientific fields separately, substantial heterogeneity emerged. Most fields showed no major gender differences, but some biomedical fields reported higher retraction rates among men, whereas fields such as engineering, economics, and information/communication technologies showed higher rates among women.

Retractions may be shaped by a complex set of factors (13–15). Beyond fraudulent or erroneous work, likelihood of retraction may depend on how intensely a paper is scrutinized and whether editorial bias affects decisions. Authors with more publications face a higher chance of retraction. This likely explains to a large extent why highly cited authors, who publish more, have higher retraction rates than non-highly cited ones, who publish less and draw less scrutiny. We cannot exclude, however, that high citations may also draw attention and thus independently increase modestly the retraction risk. Publication volume may also explain most of the small gender difference overall, as women publish slightly fewer papers on average. Higher retraction rates in non-high-income countries may reflect lower quality, higher output among top authors, potential editorial bias or a lower threshold for retracting papers. Fraudulent practices such as paper mills (16–19), cartels (20–22), extreme publishing behavior (23) and precocious citation impact (24) may also be more prevalent in these countries. Local incentive structures could drive these patterns (25,26). Whether such incentives disproportionately affect men is unclear. It may be that such perverse incentives exist more frequently in countries where there is still a large gender difference disfavoring women from publishing and reaching highly cited status.

The gender differences observed in some fields may also have complex roots. Fields where women have higher retraction rates (engineering, economics, mathematics, information & communication technologies) tend to be those where women have historically had limited presence, especially among highly cited authors, and where gender differences in authorship remain large (2). Conversely, fields where women now have lower retraction rates (biology, biomedical science) are fields where gender differences have largely disappeared. Bias or gender disadvantages may exist across diverse microenvironments in specific fields.

Younger cohorts of highly cited scientists show a substantial gender difference in retraction rates, with much higher rates in men in unadjusted analyses. However, across many fields, data for the post-2011 cohort are too sparse for meaningful interpretation and adjusted analyses did not show such a clear signal.

Among all authors, both unadjusted and adjusted analyses suggest a slightly lower risk in women in the post-2011 cohort. Any difference, if present, might appear driven by fields such as clinical medicine, biomedical research, and information & communication technologies. in clinical medicine and biomedical research, women are now better represented, especially among younger cohorts. In contrast, Information & Communication Technologies remains male-dominated. Its fast pace and competitiveness may increase risks of error or misconduct (27).

Overall, author country appears to be a much stronger correlate of retractions than gender. Prior work (3,28) highlighted the importance of country in retraction rates, with particularly high rates observed in countries that have experienced large increases (29,30)in scientific productivity over the last 20 years (29,30), such as China, India, Iran, Pakistan, Saudi Arabia, and other non-high-income countries. Our gender analysis indicates that in several of these countries, retraction rates are higher among men, particularly within the group of highly cited authors. Pakistan shows the largest disparity, which persists, though less stark, among non-highly cited authors. Nevertheless, several other countries show the opposite pattern, with higher retraction rates in women than men. This group of countries includes some high-income ones such as Italy, Taiwan, and Czech Republic.

## LIMITATIONS

We must acknowledge several limitations of this work. First, a sizeable number of authors could not have their gender assigned confidently and were therefore excluded, a situation more common in non-high-income countries. However, these gender assignment issues are unlikely to bias the observed gender–retraction association. Second, some retracted publications could not be mapped to Scopus (11), leading to underestimation of retraction proportions, although this probably also would not affect gender associations. Third, our analysis focused on large-scale entities (countries, major fields), while retraction patterns may be driven by small teams or microenvironments and occasionally tend to be clustered. In such cases, specific individuals may be primarily responsible for the retractions in such smaller units, but the role of gender, if any, in these occurrences is unclear. Fourth, we did not distinguish between honest error and misconduct. Attempting this distinction is precarious since retraction notes are often brief, vague, and inconsistently (31,32). However, most retractions appear to stem from misconduct (33). Finally, only a small number of authors among many are responsible for retractions. A recent analysis (34) involving 11,622 retracted and 19,475,437 non-retracted articles across science (Web of Science, 2008-2023) found slightly higher retraction rates (1.2-fold) for men compared to women lead authors, consistent with our findings.

## CONCLUSIONS

Our study provides a comprehensive, science-wide view of gender differences and similarities in retraction rates among highly cited and all authors, as well as the correlation of other factors with retraction rates. Gender disparities in retraction rates were overall very small or non-existent, when all scientific fields were considered. Career age, country, publication volume and scientific field were far stronger correlates of retraction risk than gender. Within specific fields, several gender differences were nevertheless noticeable in retraction risk. Fields where women have higher retraction rates tend to be those with historically limited women representation. Overall, gender appears to be a weak modulator of retraction risk, if at all, but may interact with specific fields, countries, or environments, warranting further investigation.

## ABBREVIATIONS

RWDB: Retraction Watch database
M+: Men with retractions
M-: Men without retractions
F+: Women with retractions
F-: Women without retractions

## DECLARATIONS

### Ethics approval and consent to participate

Not applicable. This study used bibliometric data and did not involve human participants.

### Consent for publication

Not applicable.

### Availability of data and materials

The grouped analytic data and analysis code used in this study are available from the corresponding author on reasonable request. The underlying Scopus data are proprietary to Elsevier. The publicly available database of top-cited scientists used for the sampling frame is available at https://elsevier.digitalcommonsdata.com/datasets/btchxktzyw/. The August 2024 dataset version was used for the analyses.

## Funding

The authors received no specific funding for this work. The work of METRICS on retractions is supported by an unrestricted gift from George F. Tidmarsh to Stanford.

### Authors’ contributions

SB conceived the study, drafted the manuscript, and supervised all stages of the project. AC contributed to study design, performed data analyses, and drafted the manuscript. JPAI contributed to study design, interpreted the results, and drafted the manuscript. AMP interpreted the results and provided supervision. JB extracted and provided data and assisted with interpretation. GR extracted and provided data, performed data analyses, and assisted with interpretation. All authors contributed to manuscript revisions and approved the final version.

## Acknowledgments

Authors wish to thank Claudia Santucci for insights into the multivariable data analysis and data visualization improvements. This work uses Scopus data provided by Elsevier.

## REFERENCES

1. De Kleijn, M, Jayabalasingham, B, Falk-Krzesinski, HJ, Collins, T, Kuiper-Hoyng, L, Cingolani, I, Zhang, J, Roberge, G, et al: The Researcher Journey Through a Gender Lens: An Examination of Research Participation, Career Progression and Perceptions Across the Globe (Elsevier, March 2020). Retrieved from https://www.elsevier.com/insights/gender-and-diversity-in-research/researcher-journey-2020. Report.

2. Ioannidis JPA, Boyack KW, Collins TA, Baas J. Gender imbalances among top-cited scientists across scientific disciplines over time through the analysis of nearly 5.8 million authors. PLoS Biol. 2023 Nov 21;21(11):e3002385. doi:10.1371/journal.pbio.3002385

3. Ioannidis JPA, Pezzullo AM, Cristiano A, Boccia S, Baas J. Linking citation and retraction data reveals the demographics of scientific retractions among highly cited authors. PLoS Biol. 2025 Jan 30;23(1):e3002999. doi:10.1371/journal.pbio.3002999

4. Pinho-Gomes AC, Hockham C, Woodward M. Women’s representation as authors of retracted papers in the biomedical sciences. PLoS One. 2023 May 3;18(5):e0284403. doi:10.1371/journal.pone.0284403

5. Oransky I. Retractions are increasing, but not enough. Nature. 2022 Aug 4;608(7921):9–9. doi:10.1038/d41586-022-02071-6

6. Archambault É, Beauchesne OH, Caruso J. Towards a Multilingual, Comprehensive and Open Scientific Journal Ontology. In. 2013. Available from: https://api.semanticscholar.org/CorpusID:85557623

7. Ioannidis JPA, Baas J, Klavans R, Boyack KW. A standardized citation metrics author database annotated for scientific field. PLoS Biol. 2019 Aug 12;17(8):e3000384. doi:10.1371/journal.pbio.3000384

8. Ioannidis JPA, Boyack KW, Baas J. Updated science-wide author databases of standardized citation indicators. PLoS Biol. 2020 Oct 16;18(10):e3000918. doi:10.1371/journal.pbio.3000918

9. NamSor. Available from: https://NamSor.app.

10. World Bank. World Bank country and lending groups. Available from: https://datahelpdesk.worldbank.org/knowledgebase/articles/906519-world-bank-country-and-lending-groups.

11. Baas J, Schotten M, Plume A, Côté G, Karimi R. Scopus as a curated, high-quality bibliometric data source for academic research in quantitative science studies. Quantitative Science Studies. 2020 Feb;1(1):377–86. doi:10.1162/qss_a_00019

12. von Elm E, Altman DG, Egger M, Pocock SJ, Gøtzsche PC, Vandenbroucke JP. The Strengthening the Reporting of Observational Studies in Epidemiology (STROBE) statement: guidelines for reporting observational studies. J Clin Epidemiol. 2008 Apr;61(4):344–9. doi:10.1016/j.jclinepi.2007.11.008

13. Steen RG, Casadevall A, Fang FC. Why has the number of scientific retractions increased? PLoS One. 2013;8(7):e68397. doi:10.1371/journal.pone.0068397 PubMed PMID: 23861902.

14. Lievore C, Rubbo P, Dos Santos CB, Picinin CT, Pilatti LA. Research ethics: a profile of retractions from world class universities. Scientometrics. 2021;126(8):6871–89. doi:10.1007/s11192-021-03987-y PubMed PMID: 34054160.

15. Li M, Shen Z. Science map of academic misconduct. Innovation (Cambridge (Mass)). 2024 Mar 4;5(2):100593. doi:10.1016/j.xinn.2024.100593 PubMed PMID: 38445017.

16. Candal-Pedreira C, Ross JS, Ruano-Ravina A, Egilman DS, Fernández E, Pérez-Ríos M. Retracted papers originating from paper mills: cross sectional study. BMJ. 2022 Nov 28;e071517. doi:10.1136/bmj-2022-071517

17. Stone R. In Iran, a shady market for papers flourishes. Science. 2016 Sep 16;353(6305):1197–1197. doi:10.1126/science.353.6305.1197

18. Abalkina A. Publication and collaboration anomalies in academic papers originating from a paper mill: Evidence from a RussiaDbased paper mill. Learned Publishing. 2023 Oct;36(4):689–702. doi:10.1002/leap.1574

19. Hvistendahl M. China’s Publication Bazaar. Science. 2013 Nov 29;342(6162):1035–9. doi:10.1126/science.342.6162.1035

20. Qiu S, Steinwender C, Azoulay P. Paper Tiger? Chinese Science and Home Bias in Citations. Cambridge, MA; 2024 May. Report. doi:10.3386/w32468

21. Plevris V. From Integrity to Inflation: Ethical and Unethical Citation Practices in Academic Publishing. J Acad Ethics. 2025 Apr 21. doi:10.1007/s10805-025-09631-1

22. Catanzaro M. Citation manipulation found to be rife in math. Science. 2024 Feb 2;383(6682):470–470. doi:10.1126/science.ado3859

23. Ioannidis JPA, Collins TA, Baas J. Evolving patterns of extreme publishing behavior across science. Scientometrics. 2024 Sep 26;129(9):5783–96. doi:10.1007/s11192-024-05117-w

24. Ioannidis JPA. Features and signals in precocious citation impact: a meta-research study. 2024. doi:10.1101/2024.10.14.618366

25. Quan W, Chen B, Shu F. Publish or impoverish. Aslib Journal of Information Management. 2017 Sep 18;69(5):486–502. doi:10.1108/AJIM-01-2017-0014

26. Abritis A, McCook A. Cash incentives for papers go global. Science. 2017 Aug 11;357(6351):541–541. doi:10.1126/science.357.6351.541

27. Memon SA, Makovi K, AlShebli B. Characterizing the effect of retractions on publishing careers. Nat Hum Behav. 2025 Jun;9(6):1134–46. doi:10.1038/s41562-025-02154-0 PubMed PMID: 40217002.

28. Sebo P, Sebo M. Geographical Disparities in Research Misconduct: Analyzing Retraction Patterns by Country. J Med Internet Res. 2025 Jan 14;27:e65775. doi:10.2196/65775

29. Thelwall M, Sud P. Scopus 1900–2020: Growth in articles, abstracts, countries, fields, and journals. Quantitative Science Studies. 2022 Apr 12;3(1):37–50. doi:10.1162/qss_a_00177

30. Bornmann L, Wagner C, Leydesdorff L. The geography of references in elite articles: Which countries contribute to the archives of knowledge? PLoS One. 2018 Mar 26;13(3):e0194805. doi:10.1371/journal.pone.0194805

31. Hwang SY, Yon DK, Lee SW, Kim MS, Kim JY, Smith L, et al. Causes for Retraction in the Biomedical Literature: A Systematic Review of Studies of Retraction Notices. J Korean Med Sci. 2023;38(41). doi:10.3346/jkms.2023.38.e333

32. Vuong Q. The limitations of retraction notices and the heroic acts of authors who correct the scholarly record: An analysis of retractions of papers published from 1975 to 2019. Learned Publishing. 2020 Apr 26;33(2):119–30. doi:10.1002/leap.1282

33. Fang FC, Steen RG, Casadevall A. Misconduct accounts for the majority of retracted scientific publications. Proceedings of the National Academy of Sciences. 2012 Oct 16;109(42):17028–33. doi:10.1073/pnas.1212247109

34. Zheng ET, Fu HZ, Thelwall M, Fang Z. Do male leading authors retract more articles than female leading authors? J Informetr. 2025 Aug;19(3):101682. doi:10.1016/j.joi.2025.101682

