## Supplementary Tables 1-6 for "Gender imbalances of retraction prevalence among highly cited authors and among all authors"

**Additional files**

**Author list:** Boccia S, Cristiano A, Pezzullo AM, Baas J, Roberge G, Ioannidis JPA

**Corresponding to:** Antonio Cristiano, MD,

**Table of contents**

**[Supplementary Table 1.](#_Toc225729117)** [Mean and median number of publications among authors across different citation groups.	2](#_Toc225729117)

**[Supplementary Table 2.](#_Toc225729118)** [Number and proportions of authors with at least one retraction across different citation groups.	3](#_Toc225729118)

**[Supplementary Table 3.](#_Toc225729119)** [Number of women and men among highly cited scientists with at least one retraction across four publication age cohorts, by field.	4](#_Toc225729119)

**[Supplementary Table 4.](#_Toc225729120)** [Number of women and men among non-highly cited scientists with at least one retraction across four publication age cohorts, by field.	5](#_Toc225729120)

**[Supplementary Table 5.](#_Toc225729121)** [Number of women and men among all scientists with at least one retraction across four publication age cohorts, by field.	6](#_Toc225729121)

**[Supplementary Table 6.](#_Toc225729122)** [Univariable and multivariable odds ratios (OR) and 95% confidence intervals (CI) from logistic regression models estimating the probability of having >=1 retraction, among all authors and top-cited authors. Publication volume is log10-transformed (number of papers). Reference categories: Men; <1992; High income level; No top-cited status; Clinical Medicine.	7](#_Toc225729122)

**[Supplementary Table 7.](#_Toc225729123)** [Completed STROBE checklist for cross-sectional studies and analysis plan information	11](#_Toc225729123)

### **Supplementary Table 1.** Mean and median number of publications among authors across different citation groups.

|  | **Highly cited** | | | | | | **Non-highly cited** | | | | | | **All citation groups** | | | | | |
| --- | --- | --- | --- | --- | --- | --- | --- | --- | --- | --- | --- | --- | --- | --- | --- | --- | --- | --- |
|  | **Men** | | | **Women** | | | **Men** | | | **Women** | | | **Men** | | | **Women** | | |
|  | **N** | **Mean** | **Median** | **N** | **Mean** | **Median** | **N** | **Mean** | **Median** | **N** | **Mean** | **Median** | **N** | **Mean** | **Median** | **N** | **Mean** | **Median** |
| **High Income Countries** |  |  |  |  |  |  |  |  |  |  |  |  |  |  |  |  |  |  |
| 1° tertile | 141230 | 205.9 | 162.0 | 27883 | 170.1 | 137.0 | 1309197 | 59.8 | 40.0 | 527023 | 49.0 | 34.0 | 1450427 | 74.0 | 42.0 | 554906 | 55.1 | 36.0 |
| 2° tertile | 420 | 15.4 | 16.0 | 118 | 15.9 | 16.0 | 1155106 | 12.9 | 12.0 | 670128 | 12.7 | 12.0 | 1155526 | 12.9 | 12.0 | 670246 | 12.7 | 12.0 |
| 3° tertile | 30 | 7.0 | 7.0 | 8 | 6.4 | 6.5 | 1094384 | 6.3 | 6.0 | 705843 | 6.3 | 6.0 | 1094414 | 6.3 | 6.0 | 705851 | 6.3 | 6.0 |
| All tertiles | 141680 | 205.3 | 162.0 | 28009 | 170.3 | 137.0 | 3558687 | 28.1 | 13.0 | 1902994 | 20.4 | 11.0 | 3700367 | 34.9 | 14.0 | 1931003 | 22.6 | 11.0 |
| **Other income levels** |  |  |  |  |  |  |  |  |  |  |  |  |  |  |  |  |  |  |
| 1° tertile | 12691 | 224.1 | 176.0 | 2975 | 235.6 | 186.0 | 447760 | 52.8 | 32.0 | 237617 | 47.2 | 34.0 | 460451 | 57.5 | 37.0 | 240592 | 49.5 | 34.0 |
| 2° tertile | 32 | 16.0 | 17 | 6 | 14.7 | 15.0 | 476600 | 12.8 | 12.0 | 333365 | 12.7 | 12.0 | 476632 | 12.8 | 12.0 | 333371 | 12.7 | 12.0 |
| 3° tertile | 1 | 8.0 | 8.0 | 0 | 0.0 | 0.0 | 538430 | 6.2 | 6.0 | 428673 | 6.2 | 6.0 | 538431 | 6.2 | 6.0 | 428673 | 6.2 | 6.0 |
| All tertiles | 12724 | 223.6 | 176.0 | 2981 | 235.1 | 185.0 | 1462790 | 22.6 | 11.0 | 999655 | 18.1 | 10.0 | 1475514 | 24.4 | 11.0 | 1002636 | 18.7 | 10.0 |
| **All income levels** |  |  |  |  |  |  |  |  |  |  |  |  |  |  |  |  |  |  |
| 1° tertile | 154885 | 206.9 | 163.0 | 31008 | 177.3 | 141.0 | 1779748 | 57.9 | 39.0 | 771303 | 48.4 | 33.0 | 1934566 | 69.8 | 42.0 | 802311 | 53.4 | 35.0 |
| 2° tertile | 410 | 14.8 | 15.0 | 129 | 15.9 | 16.0 | 1671038 | 12.9 | 12.0 | 1015910 | 12.7 | 11.0 | 1671509 | 12.9 | 12.0 | 1016039 | 12.7 | 12.0 |
| 3° tertile | 26 | 6.7 | 7.0 | 8 | 6.4 | 6.5 | 1689822 | 6.3 | 6.0 | 1153601 | 6.3 | 6.0 | 1689854 | 6.3 | 6.0 | 1153609 | 6.3 | 6.0 |
| All tertiles | 155321 | 206.4 | 162.0 | 31145 | 176.5 | 140.0 | 5140608 | 26.3 | 13.0 | 2940814 | 19.6 | 10.0 | 5295929 | 31.6 | 13.0 | 2971959 | 21.2 | 11.0 |

Authors were grouped into tertiles based on their total number of publications, with the first tertile including those with the highest papers published and the third tertile those with the least. The number of publications in the three tertiles were 20-4084, 9-19, and 5-8.

### **Supplementary Table 2.** Number and proportions of authors with at least one retraction across different citation groups.

|  | **Highly cited** | |  | **Non-highly cited** | |  | **All citation groups** | |  |
| --- | --- | --- | --- | --- | --- | --- | --- | --- | --- |
|  | **Men** | **Women** | **All** | **Men** | **Women** | **All** | **Men** | **Women** | **All** |
| **High income countries** |  |  |  |  |  |  |  |  |  |
| 1° tertile | 3865 (2.7%) | 696 (2.5%) | 5438 (2.8%) | 10397 (0.8%) | 4282(0.8%) | 17042(0.8%) | 14262 (1.0%) | 4978 (0.9%) | 22480 (1.0%) |
| 2° tertile | 0 (0.0%) | 0 (0.0%) | 1 (0.2%) | 2213 (0.2%) | 1714 (0.3%) | 4505 (0.2%) | 2213 (0.2%) | 1714 (0.3%) | 4506 (0.2%) |
| 3° tertile | 0 (0.0%) | 0 (0.0%) | 0 (0.0%) | 728 (0.1%) | 563(0.1%) | 1483 (0.1%) | 728 (0.1%) | 563 (0.1%) | 1483 (0.1%) |
| All tertiles | 3865 (2.7%) | 696 (2.5%) | 5439 (2.8%) | 13338 (0.4%) | 6559 (0.3%) | 23030 (0.4%) | 17203 (0.5%) | 7255 (0.4%) | 28469 (0.4%) |
| **Other income levels** |  |  |  |  |  |  |  |  |  |
| 1° tertile | 876 (6.9%) | 198 (6.7%) | 1627 (7.4%) | 12425 (2.8%) | 6942 (2.9%) | 28746 (3.0%) | 13301 (2.9%) | 7140 (3.0%) | 30373 (3.1%) |
| 2° tertile | 0 (0.0%) | 0 (0.0%) | 1 (2.0%) | 5419(1.2%) | 4021 (1.2%) | 14072 (1.2%) | 5419 (1.1%) | 4021 (1.2%) | 14073 (1.2%) |
| 3° tertile | 0 (0.0%) | 0 (0.0%) | 0 (0.0%) | 2517 (0.5%) | 2116 (0.5%) | 6827 (0.5%) | 2517 (0.5%) | 2116 (0.5%) | 6827 (0.4%) |
| All tertiles | 876 (6.9%) | 198 (6.6%) | 1628 (7.3%) | 20361 (1.4%) | 13079 (1.3%) | 49645 (1.4%) | 21237 (1.4%) | 13277 (1.3%) | 51273 (1.5%) |
| **All income levels** |  |  |  |  |  |  |  |  |  |
| 1° tertile | 4752 (3.1%) | 896 (2.9%) | 7081 (3.3%) | 23424 (1.3%) | 11252 (1.5%) | 45886 (1.5%) | 27631 (1.4%) | 12148 (1.5%) | 52967(1.6%) |
| 2° tertile | 0 (0.0%) | 0 (0.0%) | 2(0.3%) | 7571 (0.4%) | 5754 (0.6%) | 18653 (0.6%) | 7671 (0.5%) | 5754 (0.6%) | 18655 (0.6%) |
| 3° tertile | 0 (0.0%) | 0 (0.0%) | 0 (0.0%) | 2821 (0.2%) | 2687 (0.2%) | 8348 (0.2%) | 3266 (0.2%) | 2687 (0.2%) | 8348 (0.2%) |
| All tertiles | 4752 (3.1%) | 896 (2.9%) | 7083 (3.3%) | 33816 (0.7%) | 19693 (0.7%) | 72887 (0.7%) | 38568 (0.7%) | 20589 (0.7%) | 79970 (0.8%) |

Authors were grouped into tertiles based on their total number of publications, with the first tertile including those with the highest papers published and the third tertile those with the least.

### **Supplementary Table 3.** Number of women and men among highly cited scientists with at least one retraction across four publication age cohorts, by field.

| **Field** | **Pre-1992** | | **1992–2001** | | **2002–2011** | | **Post-2011** | |
| --- | --- | --- | --- | --- | --- | --- | --- | --- |
|  | **Women** | **Men** | **Women** | **Men** | **Women** | **Men** | **Women** | **Men** |
| Agriculture, Fisheries & Forestry | 3 (0.5%) | 22 (0.6%) | 8 (2.0%) | 25 (2.1%) | 3 (2.7%) | 12 (3.3%) | 1 (14.3%) | 1 (4.5%) |
| Biology | 18 (2.4%) | 93 (2.0%) | 14 (3.5%) | 42 (2.6%) | 0 (0.0%) | 13 (4.2%) | 1 (25.0%) | 3 (27.3%) |
| Biomedical Research | 69 (3.5%) | 487 (5.1%) | 19 (2.6%) | 103 (4.8%) | 4 (3.0%) | 18 (4.2%) | 0 (0.0%) | 1 (7.7%) |
| Built Environment & Design | 1 (1.6%) | 3 (0.8%) | 0 (0.0%) | 8 (3.0%) | 1 (2.2%) | 4 (2.3%) | 1 (33.3%) | 6 (27.3%) |
| Chemistry | 18 (2.4%) | 140 (2.1%) | 17 (2.9%) | 74 (2.9%) | 16 (5.2%) | 63 (6.0%) | 1 (8.3%) | 7 (11.1%) |
| Clinical Medicine | 255 (3.9%) | 1637 (4.5%) | 156 (4.6%) | 503 (5.4%) | 30 (3.0%) | 138 (5.5%) | 0 (0.0%) | 11 (6.5%) |
| Communication & Textual Studies | 1 (0.7%) | 0 (0.0%) | 0 (0.0%) | 0 (0.0%) | 0 (0.0%) | 1 (0.8%) | 0 (0.0%) | 0 (0.0%) |
| Earth & Environmental Sciences | 3 (0.8%) | 39 (1.0%) | 8 (2.8%) | 22 (1.9%) | 5 (4.1%) | 25 (5.8%) | 0 (0.0%) | 5 (8.2%) |
| Economics & Business | 0 (0.0%) | 14 (0.8%) | 4 (1.6%) | 16 (1.7%) | 2 (1.9%) | 13 (3.5%) | 0 (0.0%) | 1 (2.7%) |
| Enabling & Strategic Technologies | 16 (2.6%) | 112 (1.8%) | 28 (3.8%) | 125 (3.7%) | 24 (3.6%) | 120 (4.8%) | 3 (6.3%) | 27 (8.4%) |
| Engineering | 11 (2.2%) | 85 (1.2%) | 23 (4.1%) | 82 (2.4%) | 17 (4.6%) | 66 (3,5%) | 1 (3.3%) | 14 (8.9%) |
| Historical Studies | 0 (0.0%) | 0 (0.0%) | 0 (0.0%) | 0 (0.0%) | 0 (0.0%) | 0 (0.0%) | 0 (0.0%) | 0 (0.0%) |
| Information & Communication Technologies | 10 (2.1%) | 36 (0.8%) | 5 (0.8%) | 49 (1.3%) | 7 (1.6%) | 81 (3.5%) | 0 (0.0%) | 20 (9.1%) |
| Mathematics & Statistics | 2 (2.5%) | 15 (1.0%) | 1 (1.9%) | 2 (0.4%) | 3 (20.0%) | 10 (6.8%) | 0 (0.0%) | 1 (8.3%) |
| Philosophy & Theology | 0 (0.0%) | 0 (0.0%) | 0 (0.0%) | 0 (0.0%) | 0 (0.0%) | 0 (0.0%) | 0 (0.0%) | 0 (0.0%) |
| Physics & Astronomy | 17 (2.4%) | 159 (1.4%) | 5 (1.1%) | 52 (1.4%) | 6 (3.9%) | 34 (3.1%) | 2 (18.2%) | 7 (10.1%) |
| Psychology & Cognitive Sciences | 10 (1.5%) | 37 (2.1%) | 9 (3.2%) | 22 (4.0%) | 5 (5.7%) | 6 (3.3%) | 0 (0.0%) | 0 (0.0%) |
| Public Health & Health Services | 16 (1.5%) | 17 (1.6%) | 10 (1.7%) | 5 (1.1%) | 1 (0.7%) | 4 (3.1%) | 1 (25.0%) | 1 (14.3%) |
| Social Sciences | 0 (0.0%) | 6 (0.3%) | 1 (0.2%) | 6 (0,6%) | 2 (0.7%) | 1 (0.3%) | 1 (10.0%) | 0 (0.0%) |
| Visual & Performing Arts | 0 (0.0%) | 0 (0.0%) | 0 (0.0%) | 0 (0.0%) | 0 (0.0%) | 0 (0.0%) | 0 (0.0%) | 0 (0.0%) |
| All Fields | 450 (2.7%) | 2902 (2.8%) | 308 (3.0%) | 1136 (3.1%) | 126 (3.0%) | 609 (4.2%) | 12 (4.9%) | 105 (8.7%) |

### **Supplementary Table 4.** Number of women and men among non-highly cited scientists with at least one retraction across four publication age cohorts, by field.

| **Field** | **Pre-1992** | | **1992–2001** | | **2002–2011** | | **Post-2011** | |
| --- | --- | --- | --- | --- | --- | --- | --- | --- |
|  | **Women** | **Men** | **Women** | **Men** | **Women** | **Men** | **Women** | **Men** |
| Agriculture, Fisheries & Forestry | 6 (0.1%) | 49 (0.2%) | 54 (0.4%) | 128 (0.4%) | 191 (0.5%) | 433 (0.8%) | 131 (0.2%) | 217 (0.4%) |
| Biology | 27 (0.2%) | 112 (0.3%) | 87 (0.4%) | 206 (0.6%) | 264 (0.6%) | 585 (1.0%) | 189 (0.4%) | 336 (0.6%) |
| Biomedical Research | 254 (0.5%) | 672 (0.7%) | 422 (0.7%) | 743 (1.2%) | 866 (0.8%) | 1035 (1.1%) | 581 (0.5%) | 637 (0.6%) |
| Built Environment & Design | 0 (0.0%) | 4 (0.1%) | 4 (0.3%) | 16 (0.4%) | 22 (0.5%) | 51 (0.5%) | 14 (0.2%) | 40 (0.3%) |
| Chemistry | 44 (0.2%) | 197 (0.2%) | 118 (0.4%) | 280 (0.5%) | 404 (0.6%) | 649 (0.7%) | 284 (0.3%) | 432 (0.4%) |
| Clinical Medicine | 623 (0.4%) | 2111 (0.5%) | 1380 (0.9%) | 2775 (1.1%) | 4322 (1.3%) | 6391 (1.6%) | 4113 (1.0%) | 5081 (1.2%) |
| Communication & Textual Studies | 0 (0.0%) | 2 (0.0%) | 4 (0.1%) | 1 (0.0%) | 1 (0.0%) | 8 (0.1%) | 7 (0.1%) | 12 (0.2%) |
| Earth & Environmental Sciences | 9 (0.1%) | 60 (0.1%) | 53 (0.4%) | 134 (0.4%) | 269 (0.9%) | 394 (0.8%) | 133 (0.3%) | 234 (0.4%) |
| Economics & Business | 8 (0.2%) | 19 (0.1%) | 29 (0.4%) | 52 (0.3%) | 204 (1.0%) | 247 (0.7%) | 96 (0.4%) | 147 (0.4%) |
| Enabling & Strategic Technologies | 18 (0.2%) | 82 (0.1%) | 96 (0.6%) | 265 (0.4%) | 549 (1.0%) | 1011 (0.8%) | 366 (0.4%) | 882 (0.4%) |
| Engineering | 16 (0.2%) | 102 (0.1%) | 102 (0.7%) | 271 (0.4%) | 539 (1.1%) | 959 (0.7%) | 190 (0.3%) | 388 (0.2%) |
| Historical Studies | 0 (0.0%) | 8 (0.1%) | 1 (0.0%) | 5 (0.1%) | 3 (0.0%) | 8 (0.1%) | 3 (0.1%) | 7 (0.1%) |
| Information & Communication Technologies | 8 (0.2%) | 68 (0.2%) | 80 (0.7%) | 228 (0.4%) | 970 (1.8%) | 1721 (1.1%) | 500 (0.7%) | 1142 (0.6%) |
| Mathematics & Statistics | 4 (0.2%) | 32 (0.1%) | 6 (0.2%) | 41 (0.3%) | 45 (0.6%) | 115 (0.6%) | 39 (0.4%) | 120 (0.5%) |
| Philosophy & Theology | 2 (0.3%) | 1 (0.0%) | 0 (0.0%) | 2 (0.1%) | 3 (0.1%) | 5 (0.1%) | 3 (0.1%) | 5 (0.1%) |
| Physics & Astronomy | 33 (0.2%) | 239 (0.2%) | 71 (0.3%) | 270 (0.3%) | 196 (0.5%) | 463 (0.4%) | 131 (0.2%) | 354 (0.3%) |
| Psychology & Cognitive Sciences | 7 (0.1%) | 26 (0.2%) | 21 (0.2%) | 45 (0.6%) | 77 (0.4%) | 106 (0.8%) | 61 (0.2%) | 40 (0.3%) |
| Public Health & Health Services | 14 (0.1%) | 17 (0.2%) | 29 (0.2%) | 24 (0.2%) | 73 (0.2%) | 37 (0.2%) | 61 (0.1%) | 46 (0.2%) |
| Social Sciences | 5 (0.1%) | 14 (0.0%) | 10 (0.1%) | 21 (0.1%) | 38 (0.1%) | 82 (0.3%) | 109 (0.3%) | 74 (0.2%) |
| Visual & Performing Arts | 0 (0.0%) | 0 (0.0%) | 1 (0.3%) | 0 (0.0%) | 0 (0.0%) | 0 (0.0%) | 0 (0.0%) | 0 (0.0%) |
| All Fields | 1078 (0.3%) | 3815 (0.3%) | 2568 (0.6%) | 5507 (0.7%) | 9036 (1.0%) | 14300 (1.0%) | 7011 (0.6%) | 10194 (0.6%) |

### **Supplementary Table 5.** Number of women and men among all scientists with at least one retraction across four publication age cohorts, by field.

| **Field** | **Pre-1992** | | **1992–2001** | | **2002–2011** | | **Post-2011** | |
| --- | --- | --- | --- | --- | --- | --- | --- | --- |
|  | **Women** | **Men** | **Women** | **Men** | **Women** | **Men** | **Women** | **Men** |
| Agriculture, Fisheries & Forestry | 9 (0.1%) | 71 (0.2%) | 62 (0.4%) | 153 (0.5%) | 194 (0.5%) | 445 (0.9%) | 132 (0.2%) | 218 (0.4%) |
| Biology | 45 (0.4%) | 205 (0.5%) | 101 (0.5%) | 248 (0.7%) | 264 (0.6%) | 598 (1.0%) | 190 (0.4%) | 339 (0.6%) |
| Biomedical Research | 323 (0.6%) | 1159 (1.1%) | 441 (0.7%) | 846 (1.3%) | 870 (0.8%) | 1053 (1.1%) | 581 (0.5%) | 638 (0.6%) |
| Built Environment & Design | 1 (0.1%) | 7 (0.1%) | 4 (0.3%) | 24 (0.5%) | 23 (0.5%) | 55 (0.5%) | 15 (0.2%) | 46 (0.3%) |
| Chemistry | 62 (0.3%) | 337 (0.3%9 | 135 (0.5%) | 354 (0.6%) | 420 (0.7%) | 712 (0.8%) | 285 (0.3%) | 439 (0.4%) |
| Clinical Medicine | 878 (0.6%) | 3748 (0.8%) | 1536 (0.9%) | 3278 (1.2%) | 4352 (1.3%) | 6529 (1.7%) | 4113 (1.0%) | 5092 (1.2%) |
| Communication & Textual Studies | 1 (0.0%) | 2 (0.0%) | 4 (0.1%) | 1 (0.0%) | 1 (0.0%) | 9 (0.1%) | 7 (0.1%) | 12 (0.2%) |
| Earth & Environmental Sciences | 12 (0.2%) | 99 (0.2%) | 61 (0.5%) | 156 (0.5%) | 274 (1.0%) | 419 (0.9%) | 133 (0.3%) | 239 (0.4%) |
| Economics & Business | 8 (0.2%) | 33 (0.1%) | 33 (0.5%) | 68 (0.3%) | 206 (1.0%) | 260 (0.7%) | 96 (0.4%) | 148 (0.4%) |
| Enabling & Strategic Technologies | 34 (0.3%) | 194 (0.2%) | 124 (0.7%) | 390 (0.6%) | 573 (1.0%) | 1131 (0.8%) | 369 (0.4%) | 909 (0.5%) |
| Engineering | 27 (0.4%) | 187 (0.2%) | 125 (0.9%) | 353 (0.5%) | 556 (1.2%) | 1,025 (0.7%) | 191 (0.3%) | 402 (0.2%) |
| Historical Studies | 0 (0.0%) | 8 (0.1%) | 1 (0.0%) | 5 (0.1%) | 3 (0.0%) | 8 (0.1%) | 3 (0.1%) | 7 (0.1%) |
| Information & Communication Technologies | 18 (0.3%) | 104 (0.2%) | 85 (0.7%) | 277 (0.5%) | 977 (1.8%) | 1802 (1.1%) | 500 (0.7%) | 1162 (0.6%) |
| Mathematics & Statistics | 6 (0.2%) | 47 (0.2%) | 7 (0.2%) | 43 (0.3%) | 48 (0.6%) | 125 (0.6%) | 39 (0.4%) | 121 (0.5%) |
| Philosophy & Theology | 2 (0.2%) | 1 (0.0%) | 0 (0.0%) | 2 (0.1%) | 3 (0.1%) | 5 (0.1%) | 3 (0.1%) | 5 (0.1%) |
| Physics & Astronomy | 50 (0.3%) | 398 (0.3%) | 76 (0.4%) | 322 (0.4%) | 202 (0.5%) | 497 (0.4%) | 133 (0.2%) | 361 (0.3%) |
| Psychology & Cognitive Sciences | 17 (0.2%) | 63 (0.3%) | 30 (0.3%) | 67 (0.8%) | 82 (0.4%) | 112 (0.8%) | 61 (0.2%) | 40 (0.3%) |
| Public Health & Health Services | 30 (0.2%) | 34 (0.3%) | 39 (0.2%) | 29 (0.3%) | 74 (0.2%) | 41 (0.3%) | 62 (0.1%) | 47 (0.2%) |
| Social Sciences | 5 (0.0%) | 20 (0.1%) | 11 (0.1%) | 27 (0.1%) | 40 (0.1%) | 83 (0.3%) | 110 (0.3%) | 74 (0.2%) |
| Visual & Performing Arts | 0 (0.0%) | 0 (0.0%) | 1 (0.2%) | 0 (0.0%) | 0 (0.0%) | 0 (0.0%) | 0 (0.0%) | 0 (0.0%) |
| All Fields | 1528 (0.5%) | 6717 (0.5%) | 2876 (0.7%) | 6643 (0.8%) | 9162 (1.0%) | 14909 (1.0%) | 7023 (0.6%) | 10299 (0.6%) |

### **Supplementary Table 6.** Univariable and multivariable odds ratios (OR) and 95% confidence intervals (CI) from logistic regression models estimating the probability of having >=1 retraction, among all authors and top-cited authors. Publication volume is log10-transformed (number of papers). Reference categories: Men; <1992; High income level; No top-cited status; Clinical Medicine.

|  | **All authors** | | | |  | **Top-cited authors** | | | |
| --- | --- | --- | --- | --- | --- | --- | --- | --- | --- |
|  | **n** | **n retracted** | **Univariable OR (95% CI)** | **Multivariable OR (95% CI)** |  | **n** | **n retracted** | **Univariable OR (95% CI)** | **Multivariable OR (95% CI)** |
| *Gender* |  |  |  |  |  |  |  |  |  |
| Men | 5,295,929 | 38,568 | Ref | Ref |  | 155,321 | 4,752 | Ref | Ref |
| Women | 2,971,959 | 20,589 | 0.95 (0.93-0.97) | 0.92 (0.86-0.99) |  | 31,145 | 896 | 0.94 (0.87-1.01) | 1.10 (0.58-2.10) |
| Unknown | 2,093,479 | 20,813 | 1.37 (1.35-1.39) | 1.25 (1.23-1.27) |  | 30,631 | 1,435 | 1.56 (1.47-1.65) | 1.15 (1.08-1.23) |
| *Age career (year of first publication)* |  |  |  |  |  |  |  |  |  |
| <1992 | 2,306,544 | 9,911 | Ref | Ref |  | 137,364 | 4,092 | Ref | Ref |
| 1992-2001 | 1,529,389 | 12,200 | 1.86 (1.81-1.91) | 1.86 (1.81-1.92) |  | 54,909 | 1,879 | 1.15 (1.09-1.22) | 1.31 (1.23-1.39) |
| 2002-2011 | 2,889,339 | 33,469 | 2.72 (2.66-2.78) | 3.14 (3.06-3.22) |  | 23,004 | 964 | 1.42 (1.33-1.53) | 1.87 (1.72-2.02) |
| >=2012 | 3,636,095 | 24,390 | 1.56 (1.53-1.60) | 3.00 (2.91-3.09) |  | 1,820 | 148 | 2.88 (2.43-3.42) | 4.93 (4.07-5.98) |
| *Income level* |  |  |  |  |  |  |  |  |  |
| High income level | 6,454,557 | 28,469 | Ref | Ref |  | 193,694 | 5,439 | Ref | Ref |
| All other income levels | 3,482,436 | 51,273 | 3.37 (3.32-3.42) | 4.09 (4.01-4.17) |  | 22,181 | 1,628 | 2.74 (2.59-2.90) | 2.59 (2.41-2.79) |
| Unknown | 424,374 | 228 | 0.12 (0.11-0.14) | 0.26 (0.23-0.31) |  | 1,222 | 16 | 0.46 (0.28-0.75) | 0.62 (0.36-1.07) |
| *Publication volume (log10 papers)* |  |  |  |  |  |  |  |  |  |
| Per +1 unit | 10,361,367 | 79,970 | 5.65 (5.58-5.72) | 8.66 (8.49-8.84) |  | 217,097 | 7,083 | 12.00 (11.08-13.00) | 11.43 (10.41-12.56) |
| *Top-cited status* |  |  |  |  |  |  |  |  |  |
| No | 10,144,270 | 72,887 | Ref | Ref |  | 0 | 0 |  |  |
| Yes | 217,097 | 7,083 | 4.66 (4.55-4.78) | 1.42 (1.37-1.47) |  | 217,097 | 7,083 |  |  |
| *Scientific field* |  |  |  |  |  |  |  |  |  |
| Clinical Medicine | 3,280,471 | 39,335 | Ref | Ref |  | 67,839 | 3,249 | Ref | Ref |
| Agriculture, Fisheries & Forestry | 368,413 | 1,663 | 0.37 (0.36-0.39) | 0.28 (0.27-0.30) |  | 7,265 | 99 | 0.27 (0.22-0.34) | 0.37 (0.30-0.46) |
| Biology | 391,318 | 2,489 | 0.53 (0.51-0.55) | 0.43 (0.41-0.45) |  | 8,656 | 222 | 0.52 (0.46-0.60) | 0.80 (0.69-0.93) |
| Biomedical Research | 831,462 | 7,298 | 0.73 (0.71-0.75) | 0.86 (0.83-0.88) |  | 16,898 | 846 | 1.05 (0.97-1.13) | 1.42 (1.30-1.54) |
| Built Environment & Design | 59,638 | 219 | 0.30 (0.27-0.35) | 0.29 (0.25-0.34) |  | 1,243 | 34 | 0.56 (0.40-0.79) | 0.73 (0.51-1.06) |
| Chemistry | 723,834 | 3,650 | 0.42 (0.40-0.43) | 0.30 (0.29-0.31) |  | 14,911 | 462 | 0.64 (0.58-0.70) | 0.50 (0.44-0.55) |
| Communication & Textual Studies | 54,770 | 42 | 0.06 (0.05-0.09) | 0.14 (0.10-0.20) |  | 1,074 | 2 | 0.04 (<0.01-0.15) | 0.10 (0.01-0.72) |
| Earth & Environmental Sciences | 351,435 | 1,943 | 0.46 (0.44-0.48) | 0.33 (0.31-0.34) |  | 7,388 | 157 | 0.43 (0.37-0.51) | 0.50 (0.42-0.60) |
| Economics & Business | 197,907 | 1,170 | 0.49 (0.46-0.52) | 0.54 (0.50-0.58) |  | 4,137 | 59 | 0.29 (0.22-0.37) | 0.61 (0.46-0.81) |
| Enabling & Strategic Technologies | 926,899 | 5,402 | 0.48 (0.47-0.50) | 0.30 (0.29-0.31) |  | 18,317 | 654 | 0.74 (0.68-0.80) | 0.52 (0.47-0.57) |
| Engineering | 800,260 | 4,360 | 0.45 (0.44-0.47) | 0.30 (0.29-0.31) |  | 17,118 | 432 | 0.51 (0.46-0.57) | 0.44 (0.39-0.49) |
| Historical Studies | 52,202 | 37 | 0.06 (0.04-0.08) | 0.14 (0.10-0.20) |  | 1,081 | 0 | — | — |
| Information & Communication Technologies | 757,752 | 7,671 | 0.84 (0.82-0.86) | 0.55 (0.54-0.57) |  | 15,087 | 275 | 0.37 (0.33-0.42) | 0.32 (0.28-0.37) |
| Mathematics & Statistics | 126,690 | 570 | 0.37 (0.34-0.40) | 0.32 (0.29-0.35) |  | 2,693 | 48 | 0.36 (0.27-0.48) | 0.44 (0.32-0.60) |
| Philosophy & Theology | 28,024 | 22 | 0.06 (0.04-0.10) | 0.11 (0.06-0.18) |  | 523 | 0 | — | — |
| Physics & Astronomy | 851,450 | 2,706 | 0.26 (0.25-0.27) | 0.15 (0.15-0.16) |  | 19,980 | 361 | 0.37 (0.33-0.41) | 0.31 (0.28-0.35) |
| Psychology & Cognitive Sciences | 127,443 | 529 | 0.34 (0.32-0.37) | 0.59 (0.53-0.66) |  | 3,871 | 98 | 0.52 (0.42-0.63) | 0.88 (0.70-1.12) |
| Public Health & Health Services | 194,989 | 425 | 0.18 (0.16-0.20) | 0.26 (0.23-0.30) |  | 3,841 | 65 | 0.34 (0.27-0.44) | 0.48 (0.34-0.66) |
| Social Sciences | 228,291 | 438 | 0.16 (0.14-0.17) | 0.25 (0.22-0.28) |  | 5,063 | 20 | 0.08 (0.05-0.12) | 0.23 (0.14-0.38) |
| Visual & Performing Arts | 7,277 | 1 | 0.01 (<0.01-0.08) | — |  | 112 | 0 | — | — |
| *Interaction* |  |  |  |  |  |  |  |  |  |
| Women in youngest cohort (>=2012) | 1,258,531 | 7,023 | 0.69 (0.68-0.71) | 0.90 (0.87-0.94) |  | 245 | 12 | 1.53 (0.85-2.73) | 0.59 (0.31-1.12) |
| Women in all other income levels | 1,002,636 | 13,277 | 1.87 (1.84-1.91) | 1.06 (1.02-1.10) |  | 2,981 | 198 | 2.14 (1.85-2.48) | 0.92 (0.74-1.13) |
| Women in unknown income | 38,320 | 57 | 0.19 (0.15-0.25) | 2.12 (1.57-2.87) |  | 155 | 2 | 0.39 (0.10-1.56) | 0.66 (0.14-3.04) |
| Women × publication volume | 2,971,959 | 20,589 | 1.19 (1.18-1.21) | 1.00 (0.96-1.05) |  | 31,145 | 896 | 0.98 (0.95-1.02) | 0.96 (0.73-1.25) |
| Women × top-cited | 31,145 | 896 | 3.84 (3.59-4.11) | 0.92 (0.85-1.00) |  | 31,145 | 896 |  |  |
| Women × Agriculture, Fisheries & Forestry | 117,128 | 397 | 0.43 (0.39-0.48) | 0.91 (0.81-1.02) |  | 1,102 | 15 | 0.41 (0.24-0.68) | 1.15 (0.65-2.03) |
| Women × Biology | 130,669 | 600 | 0.59 (0.54-0.64) | 0.90 (0.81-0.99) |  | 1,212 | 33 | 0.83 (0.59-1.17) | 1.35 (0.90-2.00) |
| Women × Biomedical Research | 337,535 | 2,215 | 0.84 (0.81-0.88) | 0.89 (0.84-0.94) |  | 2,855 | 92 | 0.99 (0.80-1.22) | 0.71 (0.55-0.90) |
| Women × Built Environment & Design | 14,315 | 43 | 0.39 (0.29-0.52) | 1.00 (0.72-1.40) |  | 184 | 3 | 0.49 (0.16-1.54) | 0.65 (0.19-2.19) |
| Women × Chemistry | 205,678 | 902 | 0.56 (0.53-0.60) | 1.02 (0.95-1.11) |  | 1,655 | 52 | 0.96 (0.73-1.27) | 1.17 (0.84-1.62) |
| Women × Communication & Textual Studies | 23,347 | 13 | 0.07 (0.04-0.12) | 0.72 (0.37-1.38) |  | 382 | 1 | 0.08 (0.01-0.55) | 2.25 (0.14-36.35) |
| Women × Earth & Environmental Sciences | 93,869 | 480 | 0.66 (0.60-0.72) | 1.14 (1.03-1.27) |  | 817 | 16 | 0.59 (0.36-0.97) | 1.03 (0.60-1.78) |
| Women × Economics & Business | 55,950 | 343 | 0.79 (0.71-0.88) | 1.33 (1.17-1.51) |  | 614 | 6 | 0.29 (0.13-0.65) | 0.86 (0.36-2.04) |
| Women × Enabling & Strategic Technologies | 184,361 | 1,100 | 0.77 (0.72-0.82) | 1.11 (1.03-1.19) |  | 2,077 | 71 | 1.05 (0.83-1.33) | 0.93 (0.69-1.25) |
| Women × Engineering | 137,762 | 899 | 0.84 (0.79-0.90) | 1.34 (1.24-1.45) |  | 1,461 | 52 | 1.09 (0.83-1.45) | 1.38 (0.99-1.93) |
| Women × Historical Studies | 16,977 | 7 | 0.05 (0.03-0.11) | 0.51 (0.23-1.17) |  | 219 | 0 | — | — |
| Women × Information & Communication Technologies | 147,608 | 1,580 | 1.40 (1.33-1.47) | 1.25 (1.17-1.33) |  | 1,572 | 22 | 0.42 (0.27-0.64) | 0.76 (0.48-1.20) |
| Women × Mathematics & Statistics | 23,680 | 100 | 0.54 (0.45-0.66) | 1.08 (0.86-1.34) |  | 149 | 6 | 1.24 (0.55-2.82) | 2.45 (0.99-6.01) |
| Women × Philosophy & Theology | 6,528 | 8 | 0.16 (0.08-0.32) | 1.99 (0.83-4.75) |  | 89 | 0 | — | — |
| Women × Physics & Astronomy | 135,740 | 461 | 0.43 (0.40-0.48) | 1.18 (1.06-1.31) |  | 1,327 | 30 | 0.68 (0.48-0.98) | 1.18 (0.79-1.77) |
| Women × Psychology & Cognitive Sciences | 61,073 | 190 | 0.40 (0.35-0.46) | 0.85 (0.71-1.02) |  | 1,047 | 24 | 0.69 (0.46-1.04) | 1.05 (0.65-1.70) |
| Women × Public Health & Health Services | 112,480 | 205 | 0.23 (0.20-0.27) | 0.92 (0.76-1.12) |  | 1,805 | 28 | 0.47 (0.32-0.68) | 1.16 (0.70-1.94) |
| Women × Social Sciences | 94,062 | 166 | 0.23 (0.19-0.26) | 1.11 (0.92-1.35) |  | 1,482 | 4 | 0.08 (0.03-0.21) | 0.73 (0.24-2.22) |
| Women × Visual & Performing Arts | 2,671 | 1 | 0.05 (<0.01-0.34) | — |  | 38 | 0 | — | — |

### **Supplementary Table 7.** Completed STROBE checklist for cross-sectional studies and analysis plan information

This checklist was elaborated using formal items recommended for cross-sectional studies from STROBE statement (https://www.strobe-statement.org).

|  | Item No | Recommendation | Respected? | Comments and quotes |
| --- | --- | --- | --- | --- |
| **Title and abstract** | 1 | (*a*) Indicate the study’s design with a commonly used term in the title or the abstract | Yes | Study design is indicated in the Methods section of the abstract.  “We conducted a descriptive cross-sectional bibliometric analysis of a Scopus-based authors database”, Page 2 L26 |
|  |  | (*b*) Provide in the abstract an informative and balanced summary of what was done and what was found | Yes | This information is stated in the study abstract (study objective described, method and results described) |
| Introduction | | |  |  |
| Background/rationale | 2 | Explain the scientific background and rationale for the investigation being reported | Yes | Rationale and existing literature are stated in the background section |
| Objectives | 3 | State specific objectives, including any prespecified hypotheses | Yes | A statement at the end of the introduction specifies the specific goals and objectives.  “In this study, we evaluated gender distribution in retractions among highly cited and all authors worldwide with at least 5 publications, using comprehensive publication and citation data from Scopus and Retraction Watch databases, and tested differences across countries and scientific subfields.” Page 3 L68 |
| Methods | | |  |  |
| Study design | 4 | Present key elements of study design early in the paper | Yes | Study design is stated in the methods section. Key elements are all described. |
| Setting | 5 | Describe the setting, locations, and relevant dates, including periods of recruitment, exposure, follow-up, and data collection | N/A | Not This is a bibliometric study. |
| Participants | 6 | (*a*) Give the eligibility criteria, and the sources and methods of selection of participants | Yes | Study population is described is the methods section, as well as exclusion criteria.  “We focused on the highly cited scientists in the career-long ranking, including self-citations, and compared also to all Scopus authors with ≥ 5 publications” Page 5 L92 |
| Variables | 7 | Clearly define all outcomes, exposures, predictors, potential confounders, and effect modifiers. Give diagnostic criteria, if applicable | Yes | Outcome, exposure and variables used to stratify the analysis are described in the methods section. |
| Data sources/ measurement | 8* | For each variable of interest, give sources of data and details of methods of assessment (measurement). Describe comparability of assessment methods if there is more than one group | Yes | These are described in the methods section.  “We have generated a comprehensive database of the top 2% most-cited scientists in each of the 174 scientific subfields defined by the Science-Metrix classification (RRID:SCR_024471) (5). This selection was based on a composite citation index, following a methodology similar to our previous studies (6,7) […] Following a previously established approach (3), we linked Scopus author entries to the Retraction Watch database (RWDB, RRID:SCR_000654), which is the most reliable database of retractions available to date. This linkage allowed us to track the number of retractions associated with each author ID in the Scopus database.” Page 4 L74 |
| Bias | 9 | Describe any efforts to address potential sources of bias | Yes | These are described in the methods section. “Expressions of concern and corrections without retraction, retraction with republication, and retractions where it is explicit that they are due to publisher/journal error rather than author error were excluded.” Page 4 L83.  “We retained only gender assignments with a confidence score above 85%.” Page 4 L90 |
| Study size | 10 | Explain how the study size was arrived at | Yes | The methods section describes how authors were selected. |
| Quantitative variables | 11 | Explain how quantitative variables were handled in the analyses. If applicable, describe which groupings were chosen and why | Yes | These are described in the methods section.  “We categorized authors into four cohorts based on the year of their first publication year:  - Pre-1992  - 1992–2001  - 2002–2011  - Post-2011” Page 5 L93  “In each age cohort, we classified authors based on their scientific field, using the 20 major fields defined by the Science-Metrix classification” Page 5 L99  “We categorized authors by countries of residence, grouping into high-income and other incomes according to the public data from the World Bank (9)” Page 5 L100 |
| Statistical methods | 12 | (*a*) Describe all statistical methods, including those used to control for confounding | Yes | These are described in the methods section.  “We examined whether the proportion of retracted authors differed by gender within each subgroup and across different scientific domains.” Page 5 L107  “We also calculated the relative propensity R of women versus men to have a retraction among top 2% most-cited scientists and among all authors in each subfield.” Page 5 L108  “We stratified authors into tertiles based on their total number of publications, comparing patterns across low, middle, and high publishing groups. We examined the mean and median number of publications in each tertile and across all tertiles and then assessed whether differences in retraction rates varied by publication volume.” Page 5 L113 |
|  |  | (*b*) Describe any methods used to examine subgroups and interactions | N/A | Non applicable. |
|  |  | (*c*) Explain how missing data were addressed | N/A | Non applicable. |
|  |  | (*d*) If applicable, describe analytical methods taking account of sampling strategy | N/A | Non applicable. |
|  |  | (*e*) Describe any sensitivity analyses | N/A | Non applicable. |
| Results | | |  |  |
| Participants | 13* | (a) Report numbers of individuals at each stage of study—eg numbers potentially eligible, examined for eligibility, confirmed eligible, included in the study, completing follow-up, and analysed | Yes | This is described at the beginning of result section.  “Out of 10,361,367 authors with at least five full publications, 8,267,888 could be classified for gender (5,295,929 men and 2,971,959 women), while gender was uncertain for 2,093,479 (20.2%).” Page 6 L127 |
|  |  | (b) Give reasons for non-participation at each stage | N/A | Not applicable. |
|  |  | (c) Consider use of a flow diagram | No | Use of a flow diagram was not deemed appropriate |
| Descriptive data | 14* | (a) Give characteristics of study participants (eg demographic, clinical, social) and information on exposures and potential confounders | Yes | Table 1 describes authors with at least one retraction across countries of different income levels.  Table 2 describes authors with at least one retraction by subfield. |
|  |  | (b) Indicate number of participants with missing data for each variable of interest | N/A | Not applicable. |
| Outcome data | 15* | Report numbers of outcome events or summary measures | Yes | All numbers are reported in Tables |
| Main results | 16 | (*a*) Give unadjusted estimates and, if applicable, confounder-adjusted estimates and their precision (eg, 95% confidence interval). Make clear which confounders were adjusted for and why they were included | Yes | Adjusted estimates are reported in Supplementary Tables |
|  |  | (*b*) Report category boundaries when continuous variables were categorized | Yes | Publication volume was divided into tertiles, with cut‑off values provided in the footnotes of the relevant supplementary tables. |
|  |  | (*c*) If relevant, consider translating estimates of relative risk into absolute risk for a meaningful time period | N/A | Not applicable. |
| Other analyses | 17 | Report other analyses done—eg analyses of subgroups and interactions, and sensitivity analyses | N/A | Not applicable |
| Discussion | | |  |  |
| Key results | 18 | Summarise key results with reference to study objectives | Yes | Key results are presented at the beginning of the discussion section (page 10), where they are outlined in relation to the study objectives, and are further elaborated throughout the section in the context of field‑, country‑, and cohort‑specific patterns. They are also summarized in the conclusions section (page 13). |
| Limitations | 19 | Discuss limitations of the study, taking into account sources of potential bias or imprecision. Discuss both direction and magnitude of any potential bias | Yes | Description of limitations is done in the limitations section (page 12). |
| Interpretation | 20 | Give a cautious overall interpretation of results considering objectives, limitations, multiplicity of analyses, results from similar studies, and other relevant evidence | Yes | An interpretation of the findings is provided in the Discussion (pages 10–12), considering the study objectives, multiplicity of analyses, and comparisons with similar studies and relevant evidence. |
| Generalisability | 21 | Discuss the generalisability (external validity) of the study results | Yes | Given that the analysis encompassed the entire target population, the results are discussed in the context of their relevance at a global, science-wide level. |
| Other information | | |  |  |
| Funding | 22 | Give the source of funding and the role of the funders for the present study and, if applicable, for the original study on which the present article is based | Yes | Funding sources are reported in the Declarations section, under the Funding subhead (page 14) |

*Give information separately for exposed and unexposed groups.

**Note:** An Explanation and Elaboration article discusses each checklist item and gives methodological background and published examples of transparent reporting. The STROBE checklist is best used in conjunction with this article (freely available on the Web sites of PLoS Medicine at http://www.plosmedicine.org/, Annals of Internal Medicine at http://www.annals.org/, and Epidemiology at http://www.epidem.com/). Information on the STROBE Initiative is available at www.strobe-statement.org.
